# Circulating neurofilament light chain as a promising biomarker of AAV-induced dorsal root ganglia toxicity in nonclinical toxicology species

**DOI:** 10.1101/2021.12.23.473896

**Authors:** Kelly A. Fader, Ingrid D. Pardo, Ramesh C. Kovi, Christopher J. Somps, Helen Hong Wang, Vishal S. Vaidya, Shashi K. Ramaiah, Madhu P. Sirivelu

## Abstract

Adeno-associated virus (AAV)-induced dorsal root ganglia (DRG) toxicity has been observed in several nonclinical species, where lesions are characterized by neuronal degeneration/necrosis, nerve fiber degeneration, and mononuclear cell infiltration. As AAV vectors become an increasingly common platform for novel therapeutics, non-invasive biomarkers are needed to better characterize and manage the risk of DRG neurotoxicity in both nonclinical and clinical studies. Based on biological relevance, reagent availability, antibody cross-reactivity, DRG protein expression, and assay performance, neurofilament light chain (NF-L) emerged as a promising biomarker candidate. Dose- and time-dependent changes in NF-L were evaluated in male Wistar Han rats and cynomolgus monkeys following intravenous or intrathecal AAV injection, respectively. NF-L profiles were then compared against microscopic DRG lesions on Day 29 post-dosing. In animals exhibiting DRG toxicity, plasma/serum NF-L was strongly associated with the severity of neuronal degeneration/necrosis and nerve fiber degeneration, with elevations beginning as early as Day 8 in rats (≥5×10^13^ vg/kg) and Day 14 in monkeys (≥3.3×10^13^ vg/dose). Consistent with the unique positioning of DRGs outside the blood-brain barrier, NF-L in cerebrospinal fluid was only weakly associated with DRG findings. In summary, circulating NF-L is a promising biomarker of AAV-induced DRG toxicity in nonclinical species.

## INTRODUCTION

Adeno-associated virus (AAV)-based vectors have emerged as one of the most frequently used approaches for gene therapy, whereby nucleic acids are introduced into host cells to replace, suppress, or repair a gene. AAV is a naturally occurring, non-enveloped virus which contains a linear single-stranded DNA genome of ~4.7 kilobases (kb) (Balakrishnan and Jayandharan, 2014). Flanked between inverted terminal repeats (ITRs), the coding sequences of the genome can be replaced by a gene of interest and strategic promoter to yield a recombinant AAV (rAAV) vector. rAAVs are ideal for gene therapy given their: i) favorable safety profile, ii) deficient replication, iii) efficient transduction which results in stable expression in non-dividing cells, and iv) a relatively low frequency of integration into the host DNA compared to other vectors (Colella et al., 2018; Naso et al., 2017). Additionally, the various AAV serotypes exhibit preferred tropism for certain tissues, which can be combined with tissue-specific promoters or enhancers to design a selective gene therapy strategy (Li and Samulski, 2020).

Despite these advantages, challenges have arisen during the development of rAAV-based gene therapy including a limited packaging capacity (< 4.7 kb), dilution of vector genomes in actively dividing cells, and rapidly evolving regulatory requirements. Additionally, four major categories of potential AAV-induced toxicity have been identified: i) immunogenicity, ii) hepatotoxicity, iii) insertional mutagenesis, and iv) dorsal root ganglia (DRG) toxicity (Bolt et al., 2021; Colella et al., 2018). AAV-mediated immunogenicity typically involves T-cell and humoral responses to the vector capsid or transgene product, which may lead to death of the transduced cells and loss of therapeutic efficacy (Ertl and High, 2017; Kuranda et al., 2018; Ronzitti et al., 2020). Hepatic necrosis and coagulopathy have been observed in non-human primates (NHPs) treated with high doses of AAV (Hinderer et al., 2018; Palazzi et al., 2021), while acute liver failure has been observed in clinical trial patients (Feldman et al., 2020). Moreover, neonatal mice are reported to develop hepatocellular carcinoma following AAV integration at the RNA imprinted and accumulated in nucleus (Rian) locus (Chandler et al., 2015; Donsante et al., 2007). However, insertional mutagenesis has not been observed in adult mice or other nonclinical species, suggesting a species- and age-dependent mechanism.

Following AAV administration, NHPs and rodents exhibit a high incidence of microscopic DRG lesions, although clinical sensory deficits are rarely observed (Bolt et al., 2021; Hinderer et al., 2018; Hordeaux et al., 2020b; Hordeaux et al., 2019; Hordeaux et al., 2018a; Hordeaux et al., 2018b; Palazzi et al., 2021; Van Alstyne et al., 2021). These dose-dependent lesions are characterized by neuronal degeneration and/or necrosis, nerve fiber degeneration, and mononuclear cell infiltration. In one study, AAV-treated piglets exhibited severe DRG lesions accompanied by neurological deficits such as ataxia and impairments in conscious proprioception (Hinderer et al., 2018). A meta-analysis of 33 AAV studies involving 256 NHPs (rhesus and cynomolgus) revealed histopathological DRG findings across: i) five different capsids, ii) five different promoters, iii) twenty different transgenes, iv) three routes of administration [intravenous, intra-cisterna magna, intrathecal], and v) two methods of vector purification [column-purified, iodixanol-purified]. Dose and age at injection were found to significantly affect the severity of neuronal lesions, while sex had no impact (Hordeaux et al., 2020a). Taken together, these studies suggest DRG toxicity in nonclinical species is driven by the AAV modality itself rather than any specific therapeutic target. Comparable to these DRG findings, neuronal degeneration/necrosis and nerve fiber degeneration have also been observed in trigeminal and autonomic ganglia from AAV-treated rats and cynomolgus monkeys, indicating various ganglia throughout the body are affected (Palazzi et al., 2021). In humans, sensory deficits (meningoradiculitis) were observed in one of two familial amyotrophic lateral sclerosis (ALS) patients following an intrathecal infusion of AAV containing anti-superoxide dismutase 1 (SOD1) microRNA (Mueller et al., 2020). Beyond this single report, the human relevance of AAV-induced ganglia toxicity is currently unknown.

At present, there are no reports of circulating biomarkers available for assessing AAV-induced DRG toxicity in either preclinical models or humans. However, blood and cerebrospinal fluid (CSF) levels of nervous system-derived proteins have been used to monitor the progression of chronic neurodegenerative diseases and acute physical neurological damage. This includes neuron-specific proteins such as neurofilament light chain (NF-L), ubiquitin carboxy-terminal hydrolase L1 (UCH-L1), and Tau, as well as astrocyte-specific proteins such as glial fibrillary acidic protein (GFAP) and S100 calcium-binding protein B (S100B). For example, CSF NF-L, UCH-L1, total Tau, and GFAP are elevated in patients with Alzheimer’s disease (Fukuyama et al., 2001; Hendricks et al., 2019; Ohrfelt et al., 2016). These four biomarkers are also elevated in CSF, serum, and plasma from patients following traumatic brain injury (TBI; e.g. sports-related concussion) (Asken et al., 2018; Korley et al., 2018; Taghdiri et al., 2019; Zetterberg et al., 2006). The reverse translational value of these neurobiomarkers in nonclinical models has also been demonstrated. Experimental models of TBI are reported to increase serum GFAP and UCH-L1 in pigs (central fluid percussion injury) and rats (controlled cortical impact), respectively (Lafrenaye et al., 2020; Liu et al., 2010). Moreover, serum NF-L was elevated in rats treated with various neurotoxicants (e.g. chloroproprionic acid, acrylamide, trimethyltin, kainic acid), as well as rodent models of Huntington’s disease and pneumococcal meningitis (Le et al., 2020; Sano et al., 2021; Soylu-Kucharz et al., 2017; Vlasakova et al., 2019).

As rAAV vectors become an increasingly common platform for novel therapeutics, non-invasive biomarkers are needed to better characterize and evaluate the risk of DRG neurotoxicity in both nonclinical and clinical studies. The objectives of this research were to: i) identify novel non-invasive biomarker(s) for assessing AAV-induced DRG lesions in nonclinical species, and ii) develop an in-house assay for quantifying this novel biomarker. We demonstrate that NF-L is a promising biomarker of AAV-induced DRG toxicity in rats and cynomolgus monkeys, where plasma/serum increases were strongly associated with the severity of neuron degeneration/necrosis and nerve fiber degeneration.

## MATERIALS & METHODS

### Animal procedures

These studies were conducted in accordance with the current guidelines for animal welfare (National Research Council Guide for the Care and Use of Laboratory Animals, 2011 and Animal Welfare Act [AWA], 1966, as amended in 1970, 1976, 1985, and 1990, and the AWA implementing regulations in Title 9, Code of Federal Regulations, Chapter 1, Subchapter A, Parts 1-3). The procedures used in these studies have been reviewed and approved by Pfizer’s Institutional Animal Care and Use Committee.

A total of 24 male Wistar Han (Crl:WI[Han]) rats were acclimated to the facility for a minimum of 5 days prior to initiation of dosing and were aged 7-9 weeks at study start. Animals were singly housed in individually ventilated cages from receipt through a Biosafety Level-2 period and in suspended cages for the rest of the study duration. Rats were allocated to study groups using a computer-assisted randomization procedure based on pre-dose body weights. Dosing formulations for each group were prepared by diluting the stock test article in a vehicle consisting of 0.002% Pluronic F68 in 10 mM Phosphate, 350 mM NaCl, 2.7 mM KCl, 5% Sorbitol in sterile water (pH 7.4). Rats (n=6 per dose group) received a single bolus of an AAV9 variant vector (AAV-PHP.b) at 0, 2×10^13^, 5×10^13^, or 1×10^14^ vg/kg, administered through intravenous (IV) injection via the caudal vein of the tail. This neurotropic variant vector expressed the human survival motor neuron 1 (*SMN1*) transgene driven by the ubiquitous CBh (hybrid form of chicken Beta actin) promoter (AAV-PHP.b-CBh-SMN1) as previously described (Palazzi et al., 2021). Clinical observations were recorded 4 and 6 hours after the last animal was dosed on Day 1 and twice daily on non-dosing days. All rats were fasted overnight prior to necropsy and were euthanized on Day 29. To analyze circulating biomarkers, blood was collected in K_2_EDTA VACUETTE tubes (Greiner Bio-One, Monroe NC) on Days 4, 8, and 29 post-dosing. Blood was centrifuged at 2,500 g for 10 minutes to separate plasma.

A total of 6 male cynomolgus macaques of Mauritius origin aged ≥ 2.5 years at initiation of dosing were socially housed and acclimated to the laboratory environment. All animals selected for the study were seronegative (titer ≤ 1:5) for neutralizing antibodies against the AAV9 capsid to ensure unaltered tissue transduction. Animals were implanted with an intrathecal catheter in the lumbar area with a subcutaneous access port for dosing and CSF collection. Animals were sedated and placed in the Trendelenburg position with the head tilted down 15-30° below the pelvis for dose administration and kept in this position for approximately 15 minutes post-dose. Cynomolgus monkeys (n=2 per dose group) were administered a proprietary AAV9 vector by slow injection (over approximately 2 minutes) via intrathecal catheter port according to the following dosing regimens: i) three injections of 3.3×10^13^ vg, with the second and third injection administered approximately 6 and 24 hours following the first injection (total dose = 1.0×10^14^ vg/animal), ii) single injection of 8.7×10^13^ vg, or iii) single injection of 3.0×10^13^ vg. Dosing formulations were prepared using the same vehicle described above. Immediately following administration, the catheter was flushed with 0.5 mL artificial CSF. On dosing days, clinical observations were recorded prior to injection, ≤ 2 hours after the last animal was dosed, and at the end of the workday; on non-dosing days, clinical observations were recorded twice daily. Monkeys were fasted overnight prior to necropsy and were euthanized on Day 29 post-dosing. To analyze circulating biomarkers, blood was collected in Serum Separator Clot Activator VACUETTE tubes (Greiner Bio-One) on Pre-Dose Day 34 and Days 3, 7, 14, and 29 post-dose. Blood was centrifuged at 2,500 g for 10 minutes to separate serum. CSF was collected on Day 1 (pre-dose) and 29 (post-dose).

While only male rats and monkeys were included in these AAV studies, it has previously been shown that AAV-induced DRG toxicity affects males and females equally (Hordeaux et al., 2020a). To characterize baseline biomarker profiles in age-matched naïve animals, plasma and serum samples were collected from 18 naïve Wistar Han rats (8 females, 10 males) and 20 naïve cynomolgus monkeys (14 females, 6 males).

### Histopathology

Representative DRG samples with their respective nerve roots (dorsal, ventral, and spinal nerve roots) were collected bilaterally from the cervical, thoracic, and lumbar regions of the rats (C3, T6 and L4) and monkeys (C1, C4, T3, L3 and L5). Nerve roots were evaluated instead of the standard recommended nerves (sciatic, tibial, radial, median, sural, saphenous, peroneal) (Bolon et al., 2018) because neuronal degeneration is the primary toxicity associated with AAV vectors, while any effect on peripheral nerves is secondary (Hordeaux et al., 2020a). Samples were fixed in 10% neutral buffered formalin, embedded in paraffin, sectioned longitudinally at 5 μm thickness, and stained with hematoxylin and eosin (H&E) within 72 hours of collection, according to standard protocols. Slides were evaluated and reviewed by board-certified veterinary pathologists. Images were scanned using a Leica Aperio AT2 digital whole slide scanner and acquired using ImageScope viewing software (Leica Biosystems, Vista, CA). Microscopic findings were graded on the following scale: 1 = minimal, 2 = mild, 3 = moderate, 4 = marked, and 5 = severe, where the reported severity grade per animal was based on the most severe lesion observed.

### Identification of cell-specific candidate biomarkers

Proteins commonly used to evaluate cell- and structure-specific neurotoxicity via immunohistochemistry were identified as candidate biomarkers for DRG toxicity. This included: i) UCH-L1 for neurons (Wilson et al., 1988), ii) Tau and NF-L for axons (Kanaan and Grabinski, 2021; Mages et al., 2018), iii) microtubule-associated protein 2 (MAP2) for neuron dendrites (Gumy et al., 2017), iv) GFAP for astrocytes (Yun et al., 2020), v) 2’,3’-cyclic-nucleotide 3’-phosphodiesterase (CNPase) for oligodendrocytes (Toma et al., 2007), and vi) ionized calcium binding adaptor molecule 1 (IBA1) for microglia (CNS) and macrophages (PNS) (Yun et al., 2020). The expression profile of each candidate was then examined using internal transcriptomic and proteomic tissue atlases. With the exception of IBA1, each candidate was highly enriched within nervous system tissues (e.g. cerebral cortex, spinal cord, DRGs, etc.) at both the mRNA and protein levels. Although IBA1 is expressed by all macrophages throughout the body, it remained a potential candidate as no microglia-specific markers have been identified to date. Preliminary data (internal unpublished data) indicated that serum NF-L was a particularly promising candidate, where increases were detected in cynomolgus monkeys exhibiting AAV-induced DRG toxicity.

To gauge the feasibility of developing ligand-binding assays for these biomarker candidates, the commercial availability of antibodies and recombinant protein standards was evaluated (Table 1). Meso Scale Discovery (MSD) R-PLEX kits, which include a capture and detection antibody pair along with a matched calibrator, are available for human UCH-L1 (F211O), NF-L (F217X), GFAP (F211M), and total Tau (F218D), while a mouse-specific R-PLEX kit for total Tau is also available (F228E). For IBA1, a pair of monoclonal antibodies (clone EPR16588, ab178846; clone EPR16589, ab178847) and a recombinant human protein (ab105593) are available from Abcam (Waltham, MA). Similarly, monoclonal antibodies against CNPase are available from EMD Millipore (clone 11-5B, MAB326) and Cell Signaling Technology (clone D83E10, 5664), while PhosphoSolutions offers a polyclonal antibody against MAP2 (1099-MAP2). Large quantities of high-quality recombinant proteins were not commercially available for either CNPase or MAP2, and thus calibrators for these candidates would require custom synthesis.

**Table 1:**
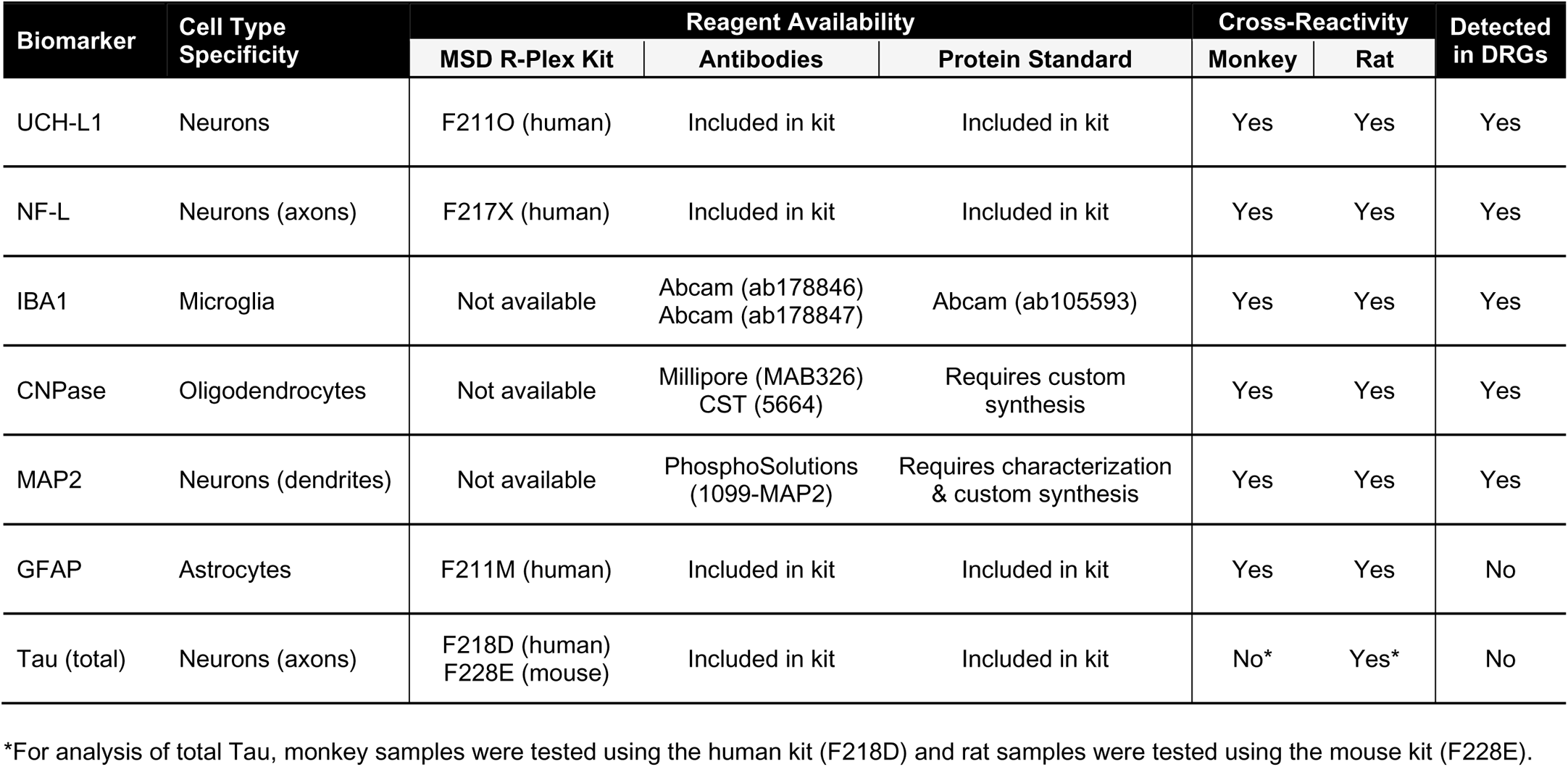
Biomarker selection and prioritization based on reagent availability, cross-reactivity, and detection in dorsal root ganglia (DRGs)

| Biomarker | Cell Type Specificity | Reagent Availability |  |  | Cross-Reactivity |  | Detected in DRGs |
| --- | --- | --- | --- | --- | --- | --- | --- |
|  |  | MSD R-Plex Kit | Antibodies | Protein Standard | Monkey | Rat |  |
| UCH-L1 | Neurons | F211O (human) | Included in kit | Included in kit | Yes | Yes | Yes |
| NF-L | Neurons (axons) | F217X (human) | Included in kit | Included in kit | Yes | Yes | Yes |
| IBA1 | Microglia | Not available | Abcam (ab178846)<br>Abcam (ab178847) | Abcam (ab105593) | Yes | Yes | Yes |
| CNPase | Oligodendrocytes | Not available | Millipore (MAB326)<br>CST (5664) | Requires custom synthesis | Yes | Yes | Yes |
| MAP2 | Neurons (dendrites) | Not available | PhosphoSolutions (1099-MAP2) | Requires characterization & custom synthesis | Yes | Yes | Yes |
| GFAP | Astrocytes | F211M (human) | Included in kit | Included in kit | Yes | Yes | No |
| Tau (total) | Neurons (axons) | F218D (human)<br>F228E (mouse) | Included in kit | Included in kit | No* | Yes* | No |

\*For analysis of total Tau, monkey samples were tested using the human kit (F218D) and rat samples were tested using the mouse kit (F228E).

### Capillary electrophoresis analysis of antibody cross-reactivity

To confirm cross-reactivity across species, each antibody discussed above was tested on brain lysate, DRG lysate, serum, and/or CSF using the ProteinSimple Wes capillary electrophoresis system (ProteinSimple, San Jose, CA). Brain and DRG samples from up to four donors per species were homogenized in Tissue Protein Extraction Reagent (T-PER) supplemented with HALT protease inhibitor and benzonase endonuclease using a Tissue Lyser Mixer Mill Grinder (Qiagen), while serum and CSF were directly diluted in supplemented T-PER. Samples were centrifuged at 10,000 g for 5 minutes and total protein in the supernatant was measured using the bicinchoninic acid (BCA) assay (Sigma). Primary antibodies were detected using a secondary antibody conjugated to horseradish peroxidase. Up to three separate plates were run for each antibody, with samples grouped as follows: i) human brain, monkey brain, monkey DRGs, ii) rat brain, rat DRGs, mouse brain, and iii) monkey serum/CSF, rat serum/CSF. Chemiluminescence signals were analyzed with Compass software (ProteinSimple), where image contrast was adjusted separately for each antibody to visualize the presence vs. absence of the target peak(s).

### MSD assay development

A duplex MSD assay was developed by combining R-PLEX kits for NF-L (F217X) and UCH-L1 (F211O) with the U-PLEX 2-Assay Development Pack (K15227N). As per the manufacturer’s protocol, biotinylated NF-L and UCH-L1 capture antibodies were incubated with U-PLEX linker #1 and #10, respectively, followed by quenching with Stop Solution. Linked capture antibodies were further diluted 10-fold in Stop Solution prior to coating the U-PLEX plate; detection antibodies were diluted 100-fold in Diluent 3. Using Diluent 43, rat plasma and serum were diluted 4-fold, while rat CSF and cynomolgus monkey serum, plasma, and CSF were diluted 6-fold. Recombinant NF-L and UCH-L1 calibrators were diluted to 25,000 and 400,000 pg/mL, respectively, in Diluent 43 to yield the top standard. An 8-point curve (including a blank) was then prepared by serial 4-fold dilutions of the top standard. The assay consisted of the following steps: i) 50 µL/well 1X duplex coating solution, ii) 50 µL/well diluted samples or standards, and iii) 50 µL/well 1X detection antibody mix. At each step, the plate was incubated with shaking (700 RPM) for 1 hour at room temperature, followed by 3 × 200 µL washes with 1X MSD Wash Buffer. Immediately following the final wash, 150 µL MSD GOLD Read Buffer A was added to each well and the plate was read on a MESO SECTOR S 600.

A ‘fit-for-purpose’ assay validation involving the following tests was performed for rat plasma: i) intra-assay precision and accuracy, ii) inter-assay precision and accuracy, iii) freeze-thaw stability, and iv) dilution linearity. Bridging validations were performed for additional biological matrices including rat CSF and serum, as well as cynomolgus monkey CSF, serum, and plasma. Only intra-assay precision/accuracy and dilution linearity were evaluated in bridging validations. Validation criteria are listed in Supplementary Table S1. Quality control (QC) samples were prepared by spiking high, mid, and low levels of NF-L and UCH-L1 calibrators into the biological matrix of interest, while detection limits were determined by diluting calibrators directly in Diluent 43. A singleplex version of the MSD assay was also developed for NF-L using the same reagents and kits described above, where spot 1 was linked to the NF-L capture antibody and spot 10 was left empty. To ensure comparability against the fully validated duplex version, a plate was run with half the wells in duplex format and the other half in singleplex format. In this side-by-side comparison, the singleplex vs. duplex values were compared for each NF-L standard on the curve, along with high, mid, and low levels of NF-L calibrator spiked into rat plasma and monkey serum.

During validation, samples and standards were run in 3 to 6 replicates, while study samples were analyzed in duplicate. For each plate, the MSD software sets the lower limit of detection (LOD) as 2.5 standard deviations above the blank. Any samples falling below the LOD were reported as the LOD value for that plate.

### Quanterix analysis of biomarkers

NF-L in rat plasma, monkey serum, and monkey CSF was quantified by Quanterix Accelerator Lab Services (Billerica, MA) using the Simoa® HD-X Neurology 4-Plex Panel B (N4PB) assay. Of the four biomarkers included in this panel, only NF-L is reported here; the assays for GFAP, UCH-L1, and total Tau exhibit poor cross-reactivity against rat and/or monkey samples and thus these data were not analyzed. Rat plasma and monkey CSF were diluted 40-fold in sample diluent, while monkey serum was diluted either 4- or 8-fold.

### Statistical analysis

Statistical analysis was performed in GraphPad Prism 9.0.0, where *p*-values ≤ 0.05 were considered significant. For biomarker data in rats, significance was evaluated through either a Kruskal-Wallis test followed by Dunn’s multiple comparison test (dose, severity) or Welch’s t-test (presence vs. absence of DRG toxicity). For biomarker data in monkeys, significance was evaluated through either one-way ANOVA analysis followed by Dunnett’s multiple comparison test (dose, severity) or t-test (presence vs. absence of DRG toxicity). For naïve biomarker levels, sex- and matrix-related differences were evaluated through two-way ANOVA analysis followed by Sidak’s multiple comparisons test. Coefficient of determination (R^2^) values were determined through linear regression analysis.

## RESULTS

### Histological DRG findings in AAV-treated rats and cynomolgus monkeys

Histopathological evaluation of H&E-stained sections of nerves attached to the DRG and neurons within the DRG revealed dose-dependent AAV-induced toxicity in male Wistar Han rats (Table 2A, Figure 1) and cynomolgus monkeys (Table 2B, Figure 2). This included minimal to moderate neuronal degeneration/necrosis, nerve fiber degeneration, and mononuclear cell infiltration in DRGs (cervical, thoracic, and lumbar regions). Nerve fiber degeneration occurred in dorsal and spinal nerve roots (sensory axons). In addition, minimal to mild satellite glial cell hypertrophy/hyperplasia was observed exclusively in rats. In the rat study, one control and one low dose (2×10^13^ vg/kg) animal exhibited minimal nerve fiber degeneration (dorsal and/or spinal nerve roots), while all other findings were observed at ≥ 5×10^13^ vg/kg. Given that minimal nerve fiber degeneration is not uncommon in rats and other laboratory animals (Pardo et al., 2020), this lesion was considered a background finding. In cynomolgus monkeys, neuronal and/or nerve fiber degeneration were observed at 3.3×10^13^ vg/dose [3x] and 8.7×10^13^ vg/dose [1x]. Despite receiving a lower total dose, one animal in the 8.7×10^13^ vg group exhibited more severe DRG lesions (moderate neuron/nerve fiber degeneration, mononuclear cell infiltration) than either animal receiving 3 injections of 3.3×10^13^ vg/dose (total = 1.0×10^14^ vg/animal).

**Figure 1:**
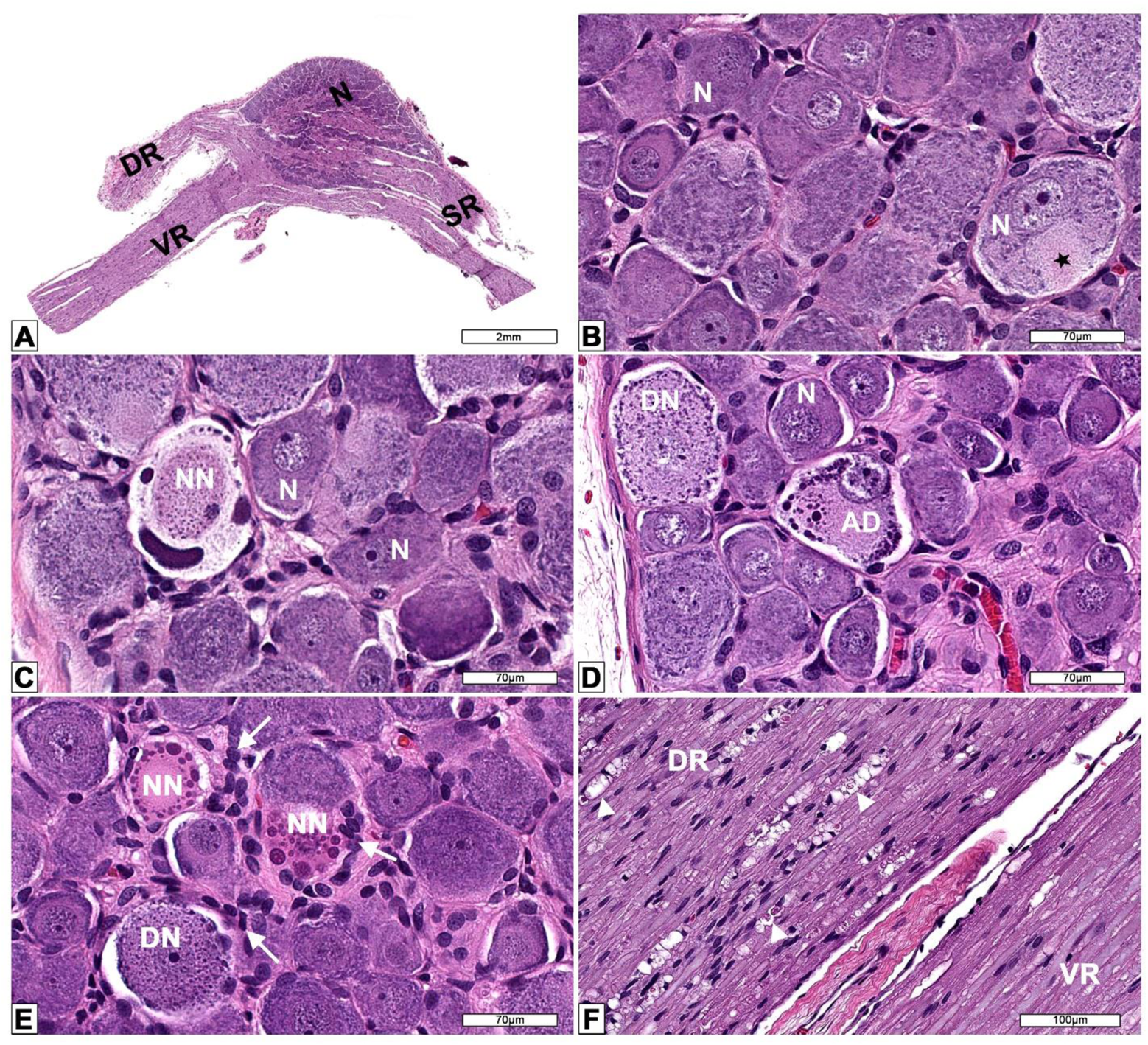
Microscopic findings in dorsal root ganglia (DRG) from AAV-treated Wistar Han rats. A) Low magnification and B) high magnification of the dorsal root ganglion (DRG) and spinal roots at lumbar 4 (L4) spinal cord segment from a male 12-week-old control Wistar Han rat. DR = dorsal spinal nerve root composed of somatic and autonomic nerve fibers (ascending sensory information passing to central nervous system). N = neurons forming the DRG and normal axon hillock (black star). SR = spinal nerve containing both sensorimotor and autonomic nerve fibers. VR = ventral spinal nerve root composed of motor and autonomic nerves (descending motor information arising from the CNS). C) DRG L4 of a male Wistar Han rat administered 1×10^14^ vg/kg of AAV showing a neuron undergoing necrosis (NN) characterized by dense basophilic round to elongated structures displaced to the periphery (nucleus and Nissl body) and central an oval eosinophilic granular material (cytoplasmic structures). D) DRG thoracic 3 from a male Wistar Han rat administered 1×10^14^ vg/kg of AAV depicting normal neurons (N) and affected neurons undergoing degeneration (DN) and another degenerated neuron showing atypical chromatolysis (AD) (cell with eccentric nucleus and eosinophilic cytoplasm with marginated Nissl bodies that formed large lumping and globular structures). E) DRG L4 from a male Wistar Han rat administered 5×10^13^ vg/kg of AAV showing reactive (hyperplastic/hypertrophic) satellite glial cells and activated macrophages (white arrows). A neuron undergoing degeneration (DN) and necrotic neurons (NN). F) Nerve roots of DRG L4 from a male Wistar Han rat administered 1×10^14^ vg/kg of AAV showing nerve fiber degeneration (white arrow heads) in the dorsal nerve root (DR) while the ventral nerve root (VR) is unremarkable. Processing: Neutral buffered 10% formalin (NBF)-fixed, paraffin embedded, hematoxylin and eosin (H&E)-stained sections.

**Figure 2:**
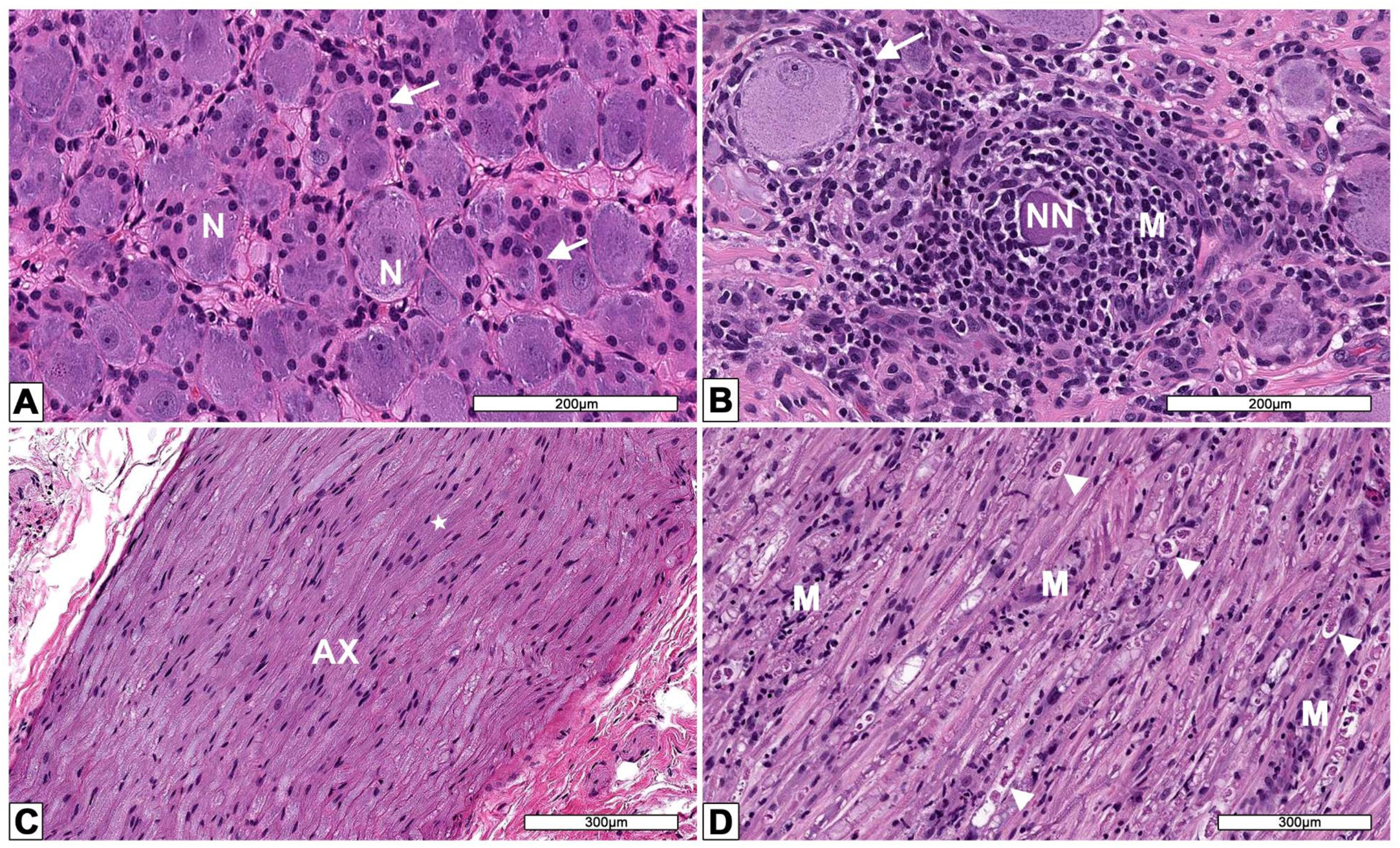
Microscopic findings in dorsal root ganglia (DRG) from AAV-treated cynomolgus monkeys. A) DRG from a male control monkey depicting healthy neurons (N) and numerous satellite glia cells (white arrows). B) DRG from a monkey administered a single injection of 8.7×10^13^ vg per animal intrathecally. A necrotic neuron (NN) is surrounded by many mononuclear cells (M). Reactive (hyperplasia/hypertrophy) satellite glial cells surround a normal neuron. C) Dorsal nerve root from a DRG of a male control monkey composed of axons (AX) surrounded by Schwann cells (white star). D) Dorsal nerve root from a DRG of a monkey administered a single injection of 8.7×10^13^ vg per animal intrathecally. Multiple arrow heads depict nerve fibers undergoing nerve fiber degeneration characterized by dilated myelin sheaths containing fragmented myelin and axons. Schwann cell reactivity and infiltration of mononuclear cells (M) are also evident. Processing: Neutral buffered 10% formalin (NBF)-fixed, paraffin embedded, hematoxylin and eosin (H&E)-stained sections.

**Table 2:** Group incidence and severity of dorsal root ganglia (DRG) microscopic findings in A) male Wistar Han rats and B) male cynomolgus monkeys.

| **Dose (vg/kg)** | **0** | **2x10<sup>13</sup>** | **5x10<sup>13</sup>** | **1x10<sup>14</sup>** |
| --- | --- | --- | --- | --- |
| **N per group** | **6** | **6** | **6** | **6** |
| ***Neuron Degeneration/Necrosis*** | 0 | 0 | 5 | 6 |
| Minimal (Grade 1) | 0 | 0 | 3 | 3 |
| Mild (Grade 2) | 0 | 0 | 2 | 3 |
| ***Nerve Fiber Degeneration*** | 1 | 1 | 5 | 6 |
| Minimal (Grade 1) | 1 | 1 | 4 | 0 |
| Mild (Grade 2) | 0 | 0 | 0 | 3 |
| Moderate (Grade 3) | 0 | 0 | 1 | 3 |
| ***Mononuclear Cell Infiltration*** | 0 | 0 | 3 | 6 |
| Minimal (Grade 1) | 0 | 0 | 1 | 6 |
| Mild (Grade 2) | 0 | 0 | 2 | 0 |
| ***Satellite Glial Cell Hypertrophy/Hyperplasia*** | 0 | 0 | 2 | 6 |
| Minimal (Grade 1) | 0 | 0 | 0 | 3 |
| Mild (Grade 2) | 0 | 0 | 2 | 3 |

**B)**
| **Dose (vg/dose)** | **3.0x10<sup>13</sup>** | **3.3x10<sup>13</sup>** | **8.7x10<sup>13</sup>** |
| --- | --- | --- | --- |
| **Number of Injections** | **1** | **3** | **1** |
| **N per group** | **2** | **2** | **2** |
| ***Neuron Degeneration/Necrosis*** | 0 | 1 | 2 |
| Minimal (Grade 1) | 0 | 0 | 0 |
| Mild (Grade 2) | 0 | 1 | 1 |
| Moderate (Grade 3) | 0 | 0 | 1 |
| ***Nerve Fiber Degeneration*** | 0 | 1 | 2 |
| Minimal (Grade 1) | 0 | 1 | 1 |
| Mild (Grade 2) | 0 | 0 | 0 |
| Moderate (Grade 3) | 0 | 0 | 1 |
| ***Mononuclear Cell Infiltration*** | 2 | 1 | 2 |
| Minimal (Grade 1) | 2 | 0 | 0 |
| Mild (Grade 2) | 0 | 1 | 1 |
| Moderate (Grade 3) | 0 | 0 | 1 |

The DRG lesions from this rat study and a similar cynomolgus monkey study have been comprehensively characterized in a previous publication, which includes regional differences in histological findings and immunohistochemical profiling of mononuclear cell infiltrates (Palazzi et al., 2021).

### Prioritization of cell-specific candidate biomarkers

To evaluate cross-reactivity against various species of interest, antibodies against each candidate biomarker were tested on brain lysate from human, cynomolgus monkey, rat, and mouse via capillary electrophoresis (ProteinSimple, Wes). Antibodies were also tested against monkey and rat DRG lysate to determine whether the candidate is present in this tissue. Both the capture and detection antibodies for UCH-L1 cross-reacted well with all four species tested. Moreover, UCH-L1 signal detected in monkey and rat DRGs was comparable to the brain (Figure 3). Interestingly, NF-L detection exhibited substantial variability both within and between species. NF-L was higher in rat DRGs compared to rat brain, while monkey NF-L was similar between the two tissues. In human and mouse brain, the NF-L detection antibody yielded a stronger signal compared to the capture antibody (Figure 3). For CNPase and IBA1, antibody cross-reactivity and presence in DRGs were confirmed for each species tested (Supplementary Figure S1A,B). GFAP was not detected in DRGs from either rat or monkey despite confirmation of cross-reactivity using brain and/or serum (Supplementary Figure S1C). Although the mouse total Tau R-PLEX antibodies exhibited good cross-reactivity with rat brain, little to no signal was detected in rat DRGs. For the human total Tau R-PLEX kit, only the capture antibody exhibited cross-reactivity with monkey brain, while neither antibody detected appreciable signal in monkey DRGs (Supplementary Figure S1D). Therefore, neither GFAP nor total Tau were considered promising candidates for monitoring DRG toxicity. Given the large molecular weight of MAP2 (~280 kDa), it is unlikely that the full-length protein would be released into circulation following moderate neurodegeneration. The presence of smaller MAP2 degradation products was therefore evaluated in serum and CSF, revealing 50 to 63 kDa fragments in both monkey and rat (Supplementary Figure S1E). Although this demonstrates feasibility for monitoring MAP2 as a non-invasive circulating biomarker, further characterization of the serum and CSF fragments would be required to synthesize an appropriate protein standard.

**Figure 3:**
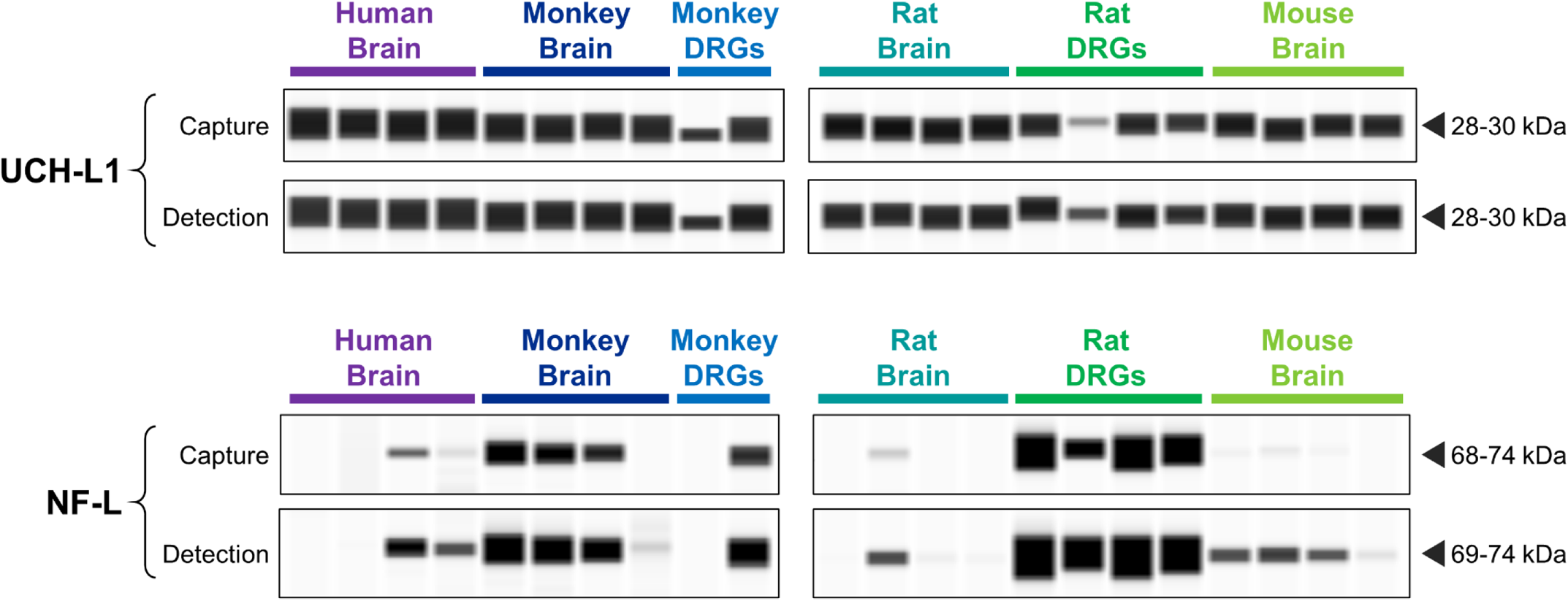
Detection of UCH-L1 and NF-L in brain and dorsal root ganglia (DRG) from nonclinical species. Wes capillary electrophoresis (ProteinSimple) was used to confirm cross-reactivity of the capture and detection antibodies within Meso Scale Discovery (MSD) R-PLEX kits for human UCH-L1 (F211O) and human NF-L (F217X). Each antibody was tested against brain homogenate (n=4) from human, cynomolgus monkey, Wistar Han rat, and CD1 mouse. Dissected DRGs from monkey (n=2) and rat (n=4) were also assessed. Chemiluminescence signals were analyzed with Compass software (ProteinSimple), where image contrast was adjusted separately for each antibody to visualize the presence vs. absence of the target peak.

Our seven candidate biomarkers were prioritized based on the factors described above. UCH-L1 and NF-L emerged as the top candidates given the availability and cross-reactivity of the MSD R-PLEX kits. CNPase, IBA1, and MAP2 were classified as a lower priority given the commercially available antibodies require conjugation to biotin and the MSD SULFO-TAG. MAP2 was further deprioritized due to the added complexity of characterizing the circulating fragments. GFAP and total Tau were excluded due to their low/undetectable levels in DRGs. The remainder of this manuscript will focus on our top biomarker candidates: NF-L and UCH-L1.

### MSD Assay Validation

Using rat plasma as the biological matrix of interest, the novel duplex MSD assay underwent a ‘fit-for-purpose’ validation, successfully passing each of the following tests: i) intra-assay accuracy and precision, ii) inter-assay accuracy and precision, iii) freeze-thaw stability, and iv) dilution linearity. More specifically, QC samples in rat plasma exhibited coefficient of variation values (CVs) ≤ 25% between replicates (intra-assay) and plates (inter-assay), where the bias between observed and expected values was within 25%. For NF-L, the analytical lower and upper limits of quantitation (LLOQ, ULOQ) were determined to be 24 and 25,000 pg/mL, while the quantitative range for UCH-L1 was determined to be 391 to 400,000 pg/mL (Supplementary Table S2). For each plate, the MSD software defines the lower limit of detection (LOD) as 2.5 standard deviations above the blank. Considering the values across each validation plate, the average analytical LOD for NF-L and UCH-L1 was 7.37 and 213 pg/mL, respectively. The functional sensitivity parameters for samples diluted 4- or 6-fold are listed in Supplementary Table S2. Compared to the initial thaw of the QC samples, 75-125% recovery was obtained following up to four freeze/thaw cycles at −80°C. Moreover, serial dilutions of 2- to 64-fold in rat plasma demonstrated linearity with a coefficient of determination (R^2^) above 0.9.

Following the full ‘fit-for-purpose’ validation in rat plasma, bridging validations were performed for additional biological matrices. Intra-assay precision and accuracy, as well as dilution linearity, were confirmed for both analytes in rat and monkey CSF. In rat serum and monkey serum/plasma, only NF-L passed the bridging validation, while the bias between observed vs. expected UCH-L1 values fell outside the acceptable limits. Specifically, observed values of UCH-L1 were more than 30% lower than expected, suggesting these matrices substantially suppress detection of this analyte. Supplementary Table S3 summarizes the validation results for each biological matrix evaluated. A bridging validation was also performed for the singleplex version of the NF-L assay. In a side-by-side comparison on a single plate, CVs between the singleplex vs. duplex versions were less than 5% for each NF-L standard and QC sample.

### NF-L in AAV-treated rats and cynomolgus monkeys

To investigate whether NF-L or UCH-L1 values were associated with DRG toxicity in rats, plasma from AAV-treated rats was analyzed using our in-house duplex MSD assay. Compared to vehicle, NF-L values were time- and dose-dependently increased in animals treated with ≥ 5×10^13^ vg/kg AAV (Figure 4A/B). In rats which exhibited DRG toxicity, NF-L was increased 9.7- and 18.7-fold on study Day 8 and 29, respectively, compared to animals with no remarkable treatment-related histological findings (Figure 4C/D). Considering the histological scores for each DRG finding, NF-L was strongly associated with the severity of both DRG neuron degeneration/necrosis (Figure 4E/F) and nerve fiber degeneration of DRG nerve roots (Figure 4G/H). In animals with mild neuronal degeneration and/or moderate nerve fiber degeneration, NF-L was increased as early as 8 days following AAV treatment, while animals with minimal neuron degeneration and/or mild nerve fiber degeneration exhibited an increase on Day 29. Focusing on Day 29, NF-L fell below the functional LLOQ (96 pg/mL) in animals with no remarkable histological findings, while all animals with mild or moderate neuron-specific findings (neuron degeneration and/or nerve fiber degeneration) exhibited values above 300 pg/mL. Mononuclear cell infiltration and glial cell hypertrophy were weakly associated with NF-L compared to neuron-specific toxicity. Interestingly, three rats exhibited elevated NF-L on Day 4. NF-L then returned to and remained at baseline by Day 8 and 29, consistent with the lack of remarkable DRG lesions in these animals (Figure 4D).

**Figure 4:**
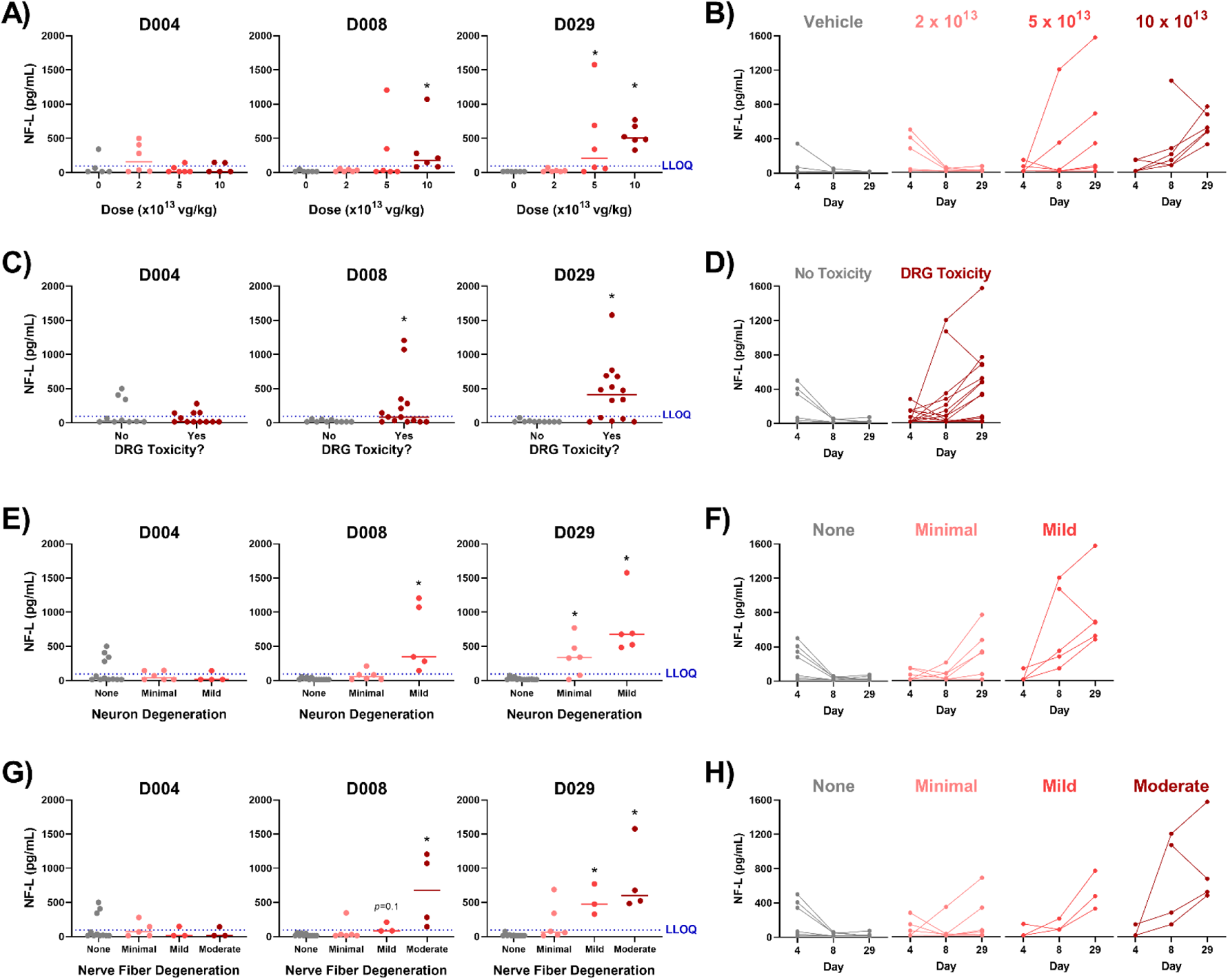
Plasma NF-L is elevated in rats exhibiting AAV-induced dorsal root ganglia (DRG) neurotoxicity. Male Wistar Han rats (n=6) were intravenously injected with a bolus dose of 0, 2×10^13^, 5×10^13^, or 1×10^14^ vg/kg AAV vector. Plasma was collected 4, 8, and 29 days after injection, while histological DRG lesions were assessed on Day 29. NF-L in plasma was quantified using an in-house duplex Meso Scale Discovery (MSD) assay. For each study day, NF-L was been plotted against A) AAV dose, C) presence/absence of DRG toxicity (neuronal and/or nerve fiber degeneration), E) severity of DRG neuron degeneration/necrosis, and G) severity of DRG nerve fiber degeneration. The dashed blue line indicates the lower limit of quantification (LLOQ = 96 pg/mL). Statistical significance (* *p* ≤ 0.05) was evaluated through a Kruskal-Wallis test followed by Dunn’s multiple comparison test (A, E, G) or Welch’s t-test (C) performed in GraphPad Prism 9.0.0. The NF-L profile for each animal has also been plotted by B) AAV dose, D) presence/absence of DRG toxicity (neuronal and/or nerve fiber degeneration), F) severity of DRG neuron degeneration/necrosis, and H) severity of DRG nerve fiber degeneration. Any values which fell below the limit of detection (LOD) were plotted as the functional LOD for that plate.

Serum samples from AAV-treated cynomolgus monkeys were also analyzed using the duplex MSD assay. On Day 29, monkeys with DRG toxicity (≥3.3×10^13^ vg/dose) exhibited a trending 4.2-fold increase in serum NF-L compared to animals with no remarkable DRG findings, with elevations beginning on Day 14 in one animal (Figure 5C/D). The lack of statistical significance is not surprising given the small number of animals used in the study. Consistent with the rat results, monkey serum NF-L was associated with the severity of DRG neuron degeneration/necrosis (Figure 5E/F) and nerve fiber degeneration of DRG nerve roots (Figure 5G/H). However, one monkey exhibited relatively high serum NF-L on Day 29 despite no remarkable DRG findings. It is possible this animal sustained procedure-related neurological injury during implantation of the intrathecal catheter, as previously reported (Boehnke et al., 2020). CSF samples from 5 of 6 monkeys on Day 29 were also analyzed; insufficient volume was available from one animal which received 3.3×10^13^ vg/dose [3x]. A monkey with mild neuron degeneration and minimal nerve fiber degeneration exhibited substantially higher NF-L in CSF compared to the other animals (Supplementary Figure S2). However, CSF NF-L from the animal with the most severe DRG toxicity (moderate neuron and nerve fiber degeneration) was comparable to animals exhibiting no DRG toxicity. Based on these results, serum and plasma were determined to be the optimal matrices for measuring NF-L as a biomarker of DRG toxicity rather than CSF.

**Figure 5:**
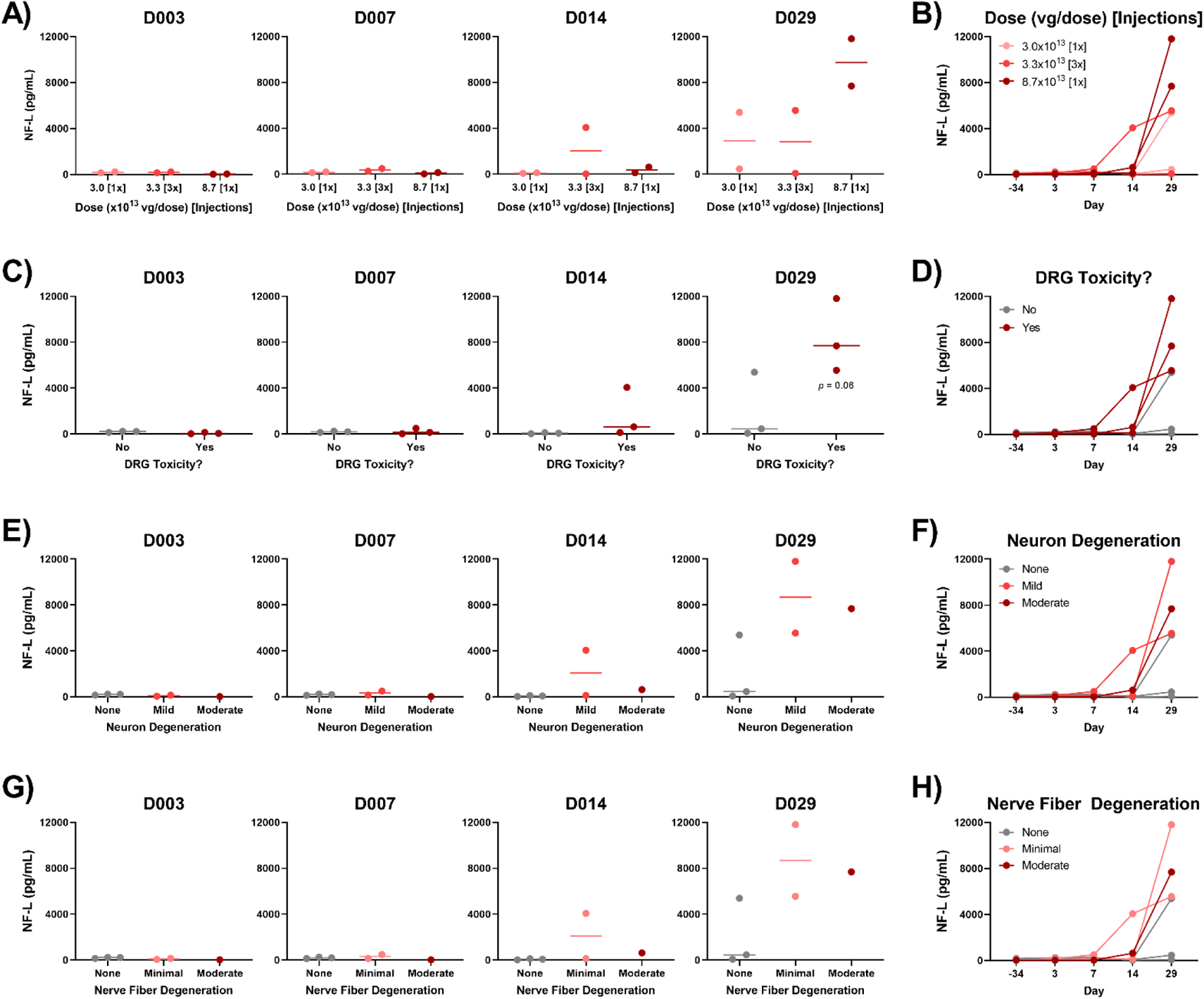
Serum NF-L is elevated in cynomolgus monkeys exhibiting AAV-induced DRG neurotoxicity. Male cynomolgus macaques (n=2) were injected with 3.0×10^13^ [1x], 3.3×10^13^ [3x], or 8.7×10^13^ [1x] vg/dose AAV vector via intrathecal catheter. Serum was collected 3, 7, 14, and 29 days after injection, while histological DRG lesions were assessed on Day 29. NF-L in serum was quantified using an in-house duplex Meso Scale Discovery (MSD) assay with a functional lower limit of quantification (LLOQ) of 144 pg/mL. For each study day, NF-L was plotted against A) AAV dose, C) presence/absence of DRG toxicity (neuronal and/or nerve fiber degeneration), E) severity of DRG neuron degeneration/necrosis, and G) severity of DRG nerve fiber degeneration. Statistical significance (*p* ≤ 0.05) was evaluated through one-way ANOVA analysis followed by Dunnett’s multiple comparison test (A, E, G) or t-test (C) performed in GraphPad Prism 9.0.0. The NF-L profile for each animal has also been plotted by B) AAV dose, D) presence/absence of DRG toxicity (neuronal and/or nerve fiber degeneration), F) severity of DRG neuron degeneration/necrosis, and H) severity of DRG nerve fiber degeneration. Any values which fell below the limit of detection (LOD) were plotted as the functional LOD for that plate.

### UCH-L1 in AAV-treated rats and cynomolgus monkeys

Throughout the study period, plasma UCH-L1 fell below the functional LLOQ (1.56 ng/mL) in all rats, with most samples also falling below the detection limit (data not shown). Similarly, UCH-L1 in serum and CSF fell below the functional LLOQ (2.35 ng/mL) in 5 of 6 monkeys, although the outliers in each matrix were not the same animal. The single monkey with quantifiable serum UCH-L1 received the lowest dose (3.0×10^13^ vg/dose [1x]) and exhibited no DRG toxicity, while the monkey with quantifiable CSF UCH-L1 exhibited mild DRG neuron degeneration with minimal nerve fiber degeneration (Supplementary Figures S3 and S4).

### Characterization of baseline NF-L in naïve animals

Given that NF-L demonstrated promise as a circulating biomarker of DRG toxicity, baseline serum and plasma profiles were characterized in age-matched naïve rats (n=18) and cynomolgus monkeys (n=20) using the singleplex version of the assay. In male rats, the average NF-L value in plasma (35.5 pg/mL) was significantly higher than in serum (24.7 pg/mL). Similarly, the average NF-L value in female rats was higher in plasma (44.1 pg/mL) compared to serum (38.2 pg/mL), although this difference failed to reach significance (*p*=0.065) (Figure 6A). All but one of the rats exhibited plasma NF-L below the functional LLOQ (96 pg/mL), with most animals falling between the LOD and LLOQ of the assay. In monkeys, serum and/or plasma NF-L fell below the LOD in 17 of the 20 animals, while only one animal exhibited values approaching the functional LLOQ (144 pg/mL) (Figure 6C). No statistically significant differences in NF-L were observed between sexes in either species. Moreover, NF-L values between plasma and serum were highly correlated in both rats and monkeys, with a coefficient of determination (R^2^) above 0.9 (Figure 6B/D).

**Figure 6:**
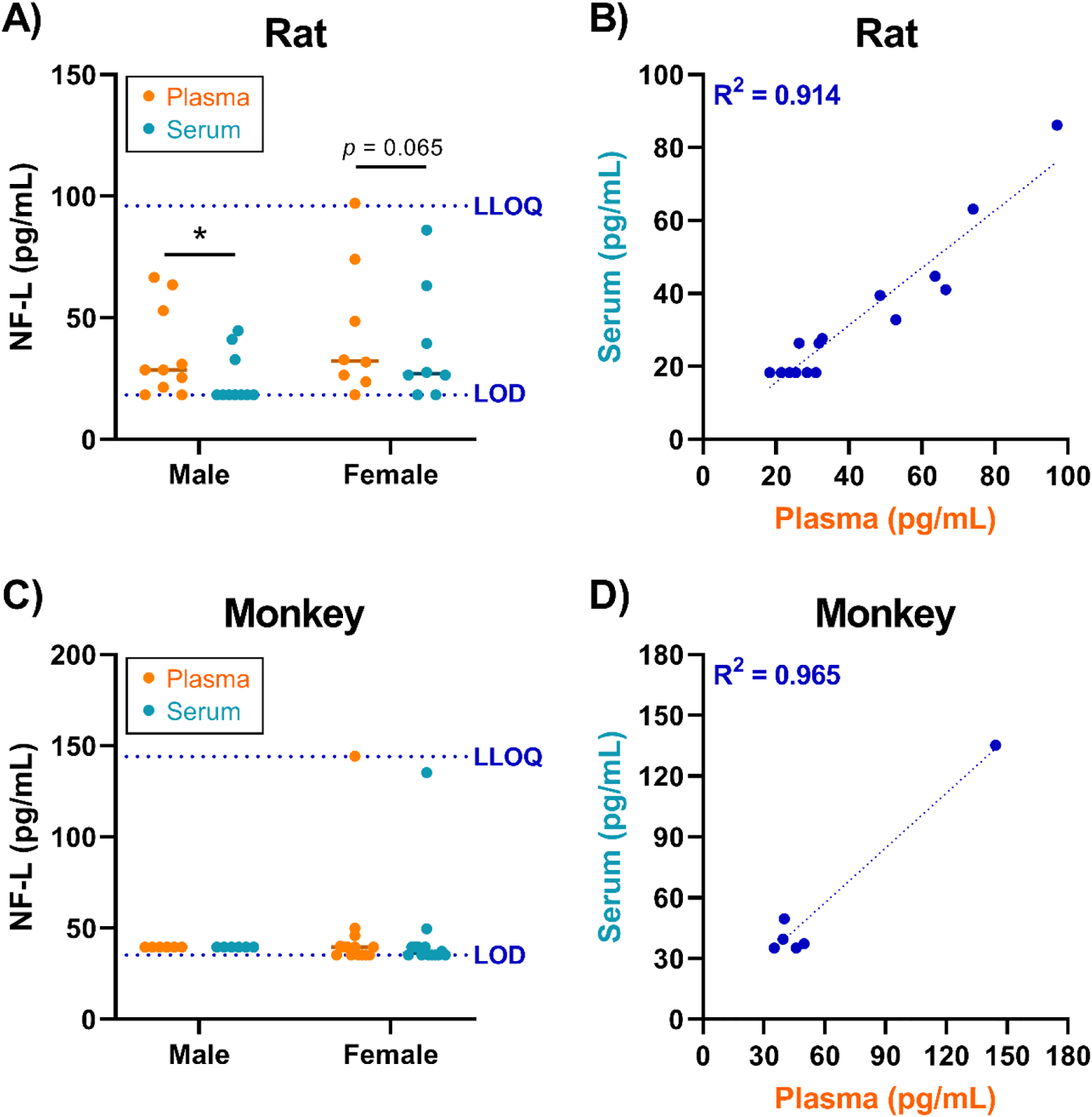
Characterization of baseline NF-L values in naïve rats and cynomolgus monkeys. An in-house singleplex Meso Scale Discovery (MSD) assay was used to quantify NF-L in matched plasma and serum from naïve A) Wistar Han rats and C) cynomolgus macaques. The dashed blue lines indicate the functional limit of detection (LOD) and lower limit of quantification (LLOQ). Statistical significance (* *p* ≤ 0.05) was evaluated through two-way ANOVA analysis followed by Sidak’s multiple comparisons test (A, C) performed in GraphPad Prism 9.0.0. The correlation between plasma and serum values of NF-L is shown for B) rats and D) monkeys, where the coefficient of determination (R^2^) was determined through linear regression analysis. Any values which fell below the limit of detection (LOD) were plotted as the functional LOD for that plate.

### MSD vs. Quanterix platform comparison

To compare the performance of our in-house MSD assay against a commercially available bead-based NF-L assay, the rat plasma and monkey serum/CSF samples from Day 29 were analyzed on the Quanterix Simoa® HD-X. The resulting Quanterix data were highly correlated with our in-house MSD values for both rat and monkey, with R^2^ values of 0.979 and 0.996, respectively (Supplementary Figure S5B/D). In rat plasma, the NF-L values determined via MSD were approximately 2-fold greater than the Quanterix values, while the MSD values for monkey serum and CSF were approximately 3-fold greater than Quanterix (Supplementary Figure S5A/C). The sensitivity of the Quanterix HD-X assay was superior to that of the MSD assay, with an analytical LLOQ of 0.5 pg/mL compared to 24 pg/mL.

## DISCUSSION

DRG toxicity observed in preclinical studies of AAV gene therapy has triggered the need for more rigorous risk assessment prior to initiation of clinical trials. During the recent meeting held by the U.S Food and Drug Administration (FDA)’s Cellular, Tissue, and Gene Therapies Advisory Committee (https://youtu.be/yLggQF0XUUY), a clear consensus emerged among panelists regarding a paucity in our current toolkit for evaluating DRG toxicity and the critical need for novel non-invasive safety biomarkers. Our studies address this key knowledge gap and help advance the DRG risk assessment strategies for AAV vectors.

In these studies, our top tier biomarker candidates (NF-L and UCH-L1) were temporally monitored in rats and cynomolgus monkeys following AAV treatment and then compared with terminal histological DRG findings on Day 29. In rats and monkeys exhibiting DRG toxicity, blood NF-L was elevated as early as Day 8 and 14, respectively. In the rat study, a separate cohort exhibited no remarkable DRG lesions 4 days after AAV treatment (Palazzi et al., 2021). A more extensive time-course study will be required to determine whether elevations in blood NF-L can be detected prior to development of overt microscopic lesions, informing the predictive value of this biomarker. Moreover, understanding whether circulating NF-L returns to baseline following recovery of DRG neurotoxicity will be important for risk assessment in humans. Reversibility of NF-L changes following recovery has previously been reported in patients with oxaliplatin-induced peripheral neuropathy, where serum levels were significantly decreased six months after completion of chemotherapy compared to levels detected during treatment (Kim et al., 2020).

Circulating NF-L in naïve animals, as well as vehicle- and AAV-treated animals lacking DRG toxicity, lie within a narrow range. The low biological variability in baseline NF-L enabled the biological sensitivity of this biomarker, where statistically significant elevations were detected in animals with low-grade histological lesions including minimal (grade 1) neuron degeneration and mild (grade 2) nerve fiber degeneration. As expected, the association between blood NF-L and severity of neuron/nerve fiber degeneration was stronger than the association with either mononuclear cell infiltration or glial cell hypertrophy. It is well known that neurofilaments such as NF-L are expressed primarily in the axon of a neuron, and thus glial cells do not contribute directly to the circulating neurofilament pool (Yuan et al., 2017). This restricted expression profile makes NF-L a highly specific marker of neuroaxonal degeneration, reducing the risk of false positives.

To our knowledge, this is the first study to demonstrate the use of NF-L as a circulating biomarker of DRG-related neurotoxicity. However, previous rodent studies report increased NF-L in experimental models of Huntington’s Disease and pneumococcal meningitis (Le et al., 2020; Soylu-Kucharz et al., 2017). Moreover, circulating NF-L was elevated in patients with TBI (Korley et al., 2018; Shahim et al., 2017; Zetterberg et al., 2006) and several neurogenerative diseases including multiple sclerosis, Alzheimer’s disease, ALS, and Guillain-Barré syndrome (Gaiottino et al., 2013; Hendricks et al., 2019; Norgren et al., 2003). In each of these studies, NF-L was elevated in CSF as well as blood-derived biofluids. Notably, in our monkey study, NF-L in CSF was weakly associated with DRG toxicity compared to serum. This is consistent with the anatomy of DRGs, which lie outside the blood-brain barrier and are surrounded by only a thin layer of CSF (Haberberger et al., 2019). Moreover, the interface between DRGs and blood vessels is highly unique, where fenestrated capillaries readily irrigate sensory neurons with a robust supply of blood to fulfill their metabolic needs (Esposito et al., 2019; Godel et al., 2016; Haberberger et al., 2019; Kiernan, 1996). Blood-borne molecules can easily enter DRG neurons, and in turn, DRG-derived molecules are released directly into circulation. This unique anatomy, combined with our comparison of monkey NF-L across matrices, suggest blood-derived biofluids (serum, plasma) are more appropriate than CSF for monitoring biomarkers of DRG toxicity. In summary, NF-L is a versatile biomarker for both central (CNS) and peripheral (PNS) neurotoxicity, although the ideal matrix for analysis may depend on the specific neurons targeted.

Prior to biomarker analysis, the presence of UCH-L1 protein in rat and monkey DRGs was confirmed through capillary electrophoresis, with levels comparable to the brain. However, regardless of AAV treatment or DRG toxicity, plasma UCH-L1 fell below the functional LLOQ in all rat plasma samples and all but one monkey serum samples analyzed. The utility of UCH-L1 as a nonclinical biomarker of AAV-induced DRG toxicity therefore remains unclear as it is possible our in-house assay simply lacks the sensitivity required to detect modest changes. Interestingly, a time-course analysis of UCH-L1 in a rat model of TBI found serum values peaked between 2 and 6 hours following cortical impact, with values approaching baseline 36 hours after injury (Liu et al., 2010). Similarly, UCH-L1 CSF from TBI patients were reported to peak 6 hours following concussion and then decrease over time (Papa et al., 2010). Following AAV treatment, the earliest timepoints evaluated in our monkey and rat studies were 72 and 96 hours, respectively, and thus evaluating serum UCH-L1 at an earlier timepoint would be worthwhile. However, a biomarker which is only transiently increased following treatment may be difficult to reliably detect, monitor, and compare across subjects. Compared to the short half-life of UCH-L1 (Kawata et al., 2016), NF-L is stable in both blood and CSF for weeks to months, allowing for flexibility in the blood collection timeline (Bergman et al., 2016).

CNPase, IBA1, and MAP2 were identified as promising candidates for evaluating the role of oligodendrocytes, macrophages/microglia, and neuronal dendrite alterations, respectively, in DRG toxicity. However, additional studies are needed to examine whether blood concentrations of these candidates correlate with AAV-elicited histological findings. It is important to note that IBA1 is expressed by all macrophages throughout the body and thus it would be difficult to distinguish microglia activation from other AAV-induced pathologies such as immunogenicity and hepatotoxicity. MAP2 has proven valuable for assessing dendrite-specific neuronal injury via immunohistochemistry (Li et al., 2000; Schartz et al., 2016; Sundaramoorthy et al., 2020). Due to its large size however, it is unlikely that the full-length version (~280 kDa) of this protein would be present in circulation. We detected small MAP2 fragments (50-63 kDa) in serum and CSF from naïve rats and cynomolgus monkeys. This is consistent with previous studies which report 50 and 70 kDa MAP2 fragments in serum from rats following ischemia/reperfusion injury (Park et al., 2012) and humans with bipolar depression (Daftary et al., 2017), respectively. The presence of small MAP2 fragments in easily accessible bodily fluids confirms the feasibility of using this candidate as a non-invasive biomarker of neuronal dendrite injury.

In conclusion, NF-L is a sensitive blood-based biomarker of DRG toxicity which can be used to non-invasively assess AAV-induced neurotoxicity in rats and cynomolgus monkeys. Moreover, the translational value of NF-L exhibits great promise, where blood values could be used to manage risk for AAV-induced DRG toxicity in human clinical trials. Notably, age-dependent reference values of serum NF-L have been determined in healthy adults (Bridel et al., 2021b; Khalil et al., 2020), while elevated serum NF-L is reported to be predictive of chemotherapy-induced peripheral neuropathy (Huehnchen et al., submitted; Kim et al., 2020). The Innovative Medicines Initiative (IMI) Translational Safety Biomarker Pipeline (TransBioLine) project includes a workstream to qualify serum NF-L, along with other nervous system-derived proteins, as biomarkers of drug-induced CNS injury in humans (Leptak and Kozauer, 2020; U.S. FDA, 2021). Although these developments are encouraging, further research is needed to understand human serum profiles of NF-L before it can be implemented in AAV clinical trials. Specifically, it will be important to determine how pre-existing neurological diseases affect detection of AAV-induced NF-L changes in patients receiving gene therapy (Bridel et al., 2021a). Once further characterized across diverse human populations, this sensitive biomarker of DRG toxicity could become a fundamental tool for monitoring the safety of novel AAV-delivered therapeutics.

## ACKNOWLEDGMENTS

The authors thank Joseph T. Brady and Jon C. Cook for their support throughout the project and review of the manuscript; Thomas A. Lanz and Chang-Ning Liu for critical review of the manuscript; June Liu for her help with study conduct and protocols; Colleen M. Doshna, Lauren Martin, and Kimberly S. Ebersole for performing QC of the study data; Ryan Criswell, Taylor Hulten, Linda M. Hutter, James A. Stejskal, and Gretchen L. Volberg for their help with sample procurement.

## AUTHOR CONTRIBUTIONS

KAF, IDP, CJS, HHW, and MPS conceptualized the studies; IDP and RCK performed histopathology assessment and review; KAF conducted method development and sample analysis; KAF and MPS analyzed and interpreted data; KAF and MPS drafted the manuscript; KAF, IDP, CJS, RCK, HHW, VSV, SKR, and MPS critically reviewed and revised the manuscript.

