## Supplementary Materials for "Circulating neurofilament light chain as a promising biomarker of AAV-induced dorsal root ganglia toxicity in nonclinical toxicology species"

Kelly A. Fader[1], Ingrid D. Pardo[2], Ramesh C. Kovi[3], Christopher J. Somps[1], Helen Hong Wang[3], Vishal S. Vaidya[3], Shashi K. Ramaiah[3], and Madhu P. Sirivelu[3]*.

[1] Pfizer Worldwide Research, Development and Medical; Drug Safety Research and Development; Groton, Connecticut

[2] Biogen; Cambridge, Massachusetts

[3] Pfizer Worldwide Research, Development and Medical; Drug Safety Research and Development; Cambridge, Massachusetts

Pfizer Inc., 300 Technology Square, Cambridge, MA, 02139, United States

#### SUPPLEMENTARY FIGURES

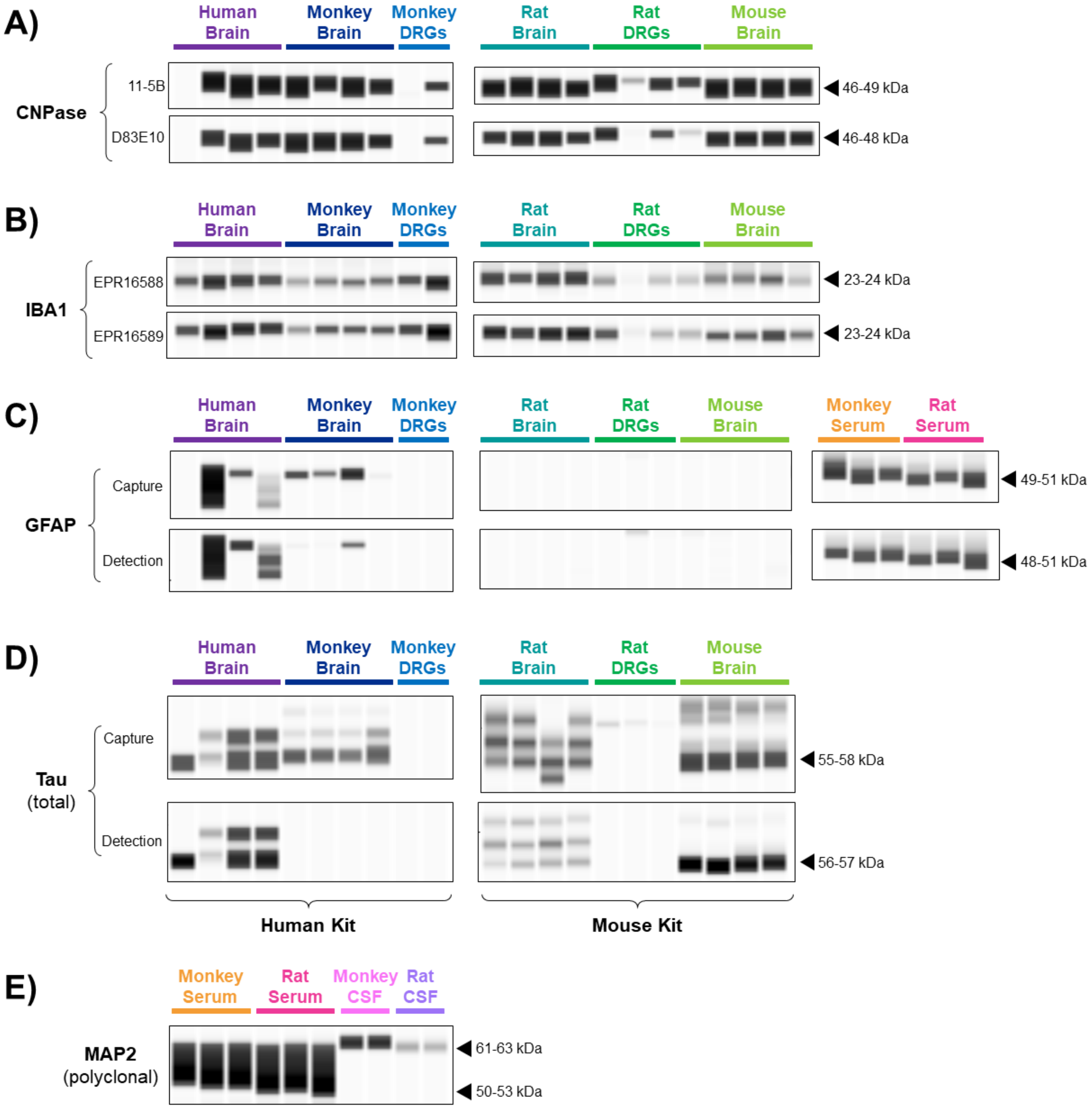


**Supplementary Figure S1: Detection of candidate biomarkers in brain, dorsal root ganglia (DRG), and biofluids from human and nonclinical species.** Wes capillary electrophoresis (ProteinSimple) was used to evaluate antibody cross-reactivity against brain homogenate (n=4) from human, cynomolgus monkey, Wistar Han rat, and CD1 mouse. Dissected DRGs (n=2-4), serum (n=3), and/or CSF (n=2) from monkey and rat were also assessed. A) CNPase antibody clones 11-5B and D83E10 were obtained from Millipore (MAB326) and Cell Signaling Technologies (5664), respectively. B) IBA1 antibody clones EPR16588 and ERP16589 were obtained from Abcam (ab178846 and ab178847, respectively). Meso Scale Discovery (MSD) R-PLEX kits were available for C) GFAP (human: F211M) and D) total Tau (human: F218D; mouse: F228E). E) The polyclonal antibody against MAP2 was obtained from PhosphoSolutions (1099-MAP2). Chemiluminescence signals were analyzed with Compass software (ProteinSimple), where image contrast was adjusted separately for each antibody to visualize the presence vs. absence of the target peak(s).

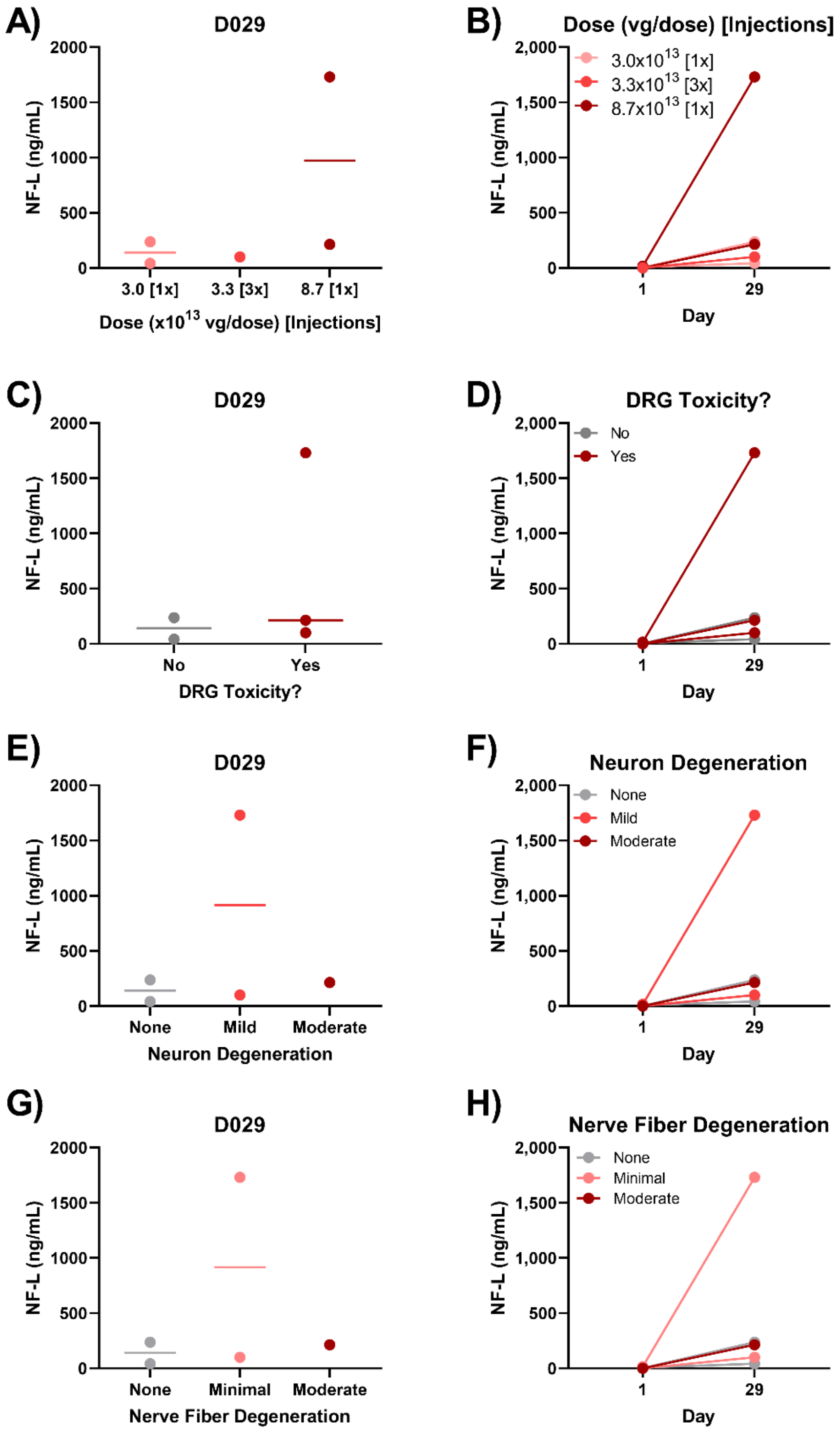


**Supplementary Figure S2: NF-L in cerebrospinal fluid (CSF) from cynomolgus monkeys exhibiting AAV-induced dorsal root ganglia (DRG) toxicity.** Male cynomolgus macaques (n=2 per group) were injected with $3.0 \times 10^{13}$ [1x], $3.3 \times 10^{13}$ [3x], or $8.7 \times 10^{13}$ [1x] vg/dose AAV vector via intrathecal catheter. CSF was collected on Day 1 (pre-dose) and 29, while histological DRG lesions were assessed on Day 29. NF-L in CSF was quantified using an in-house duplex Meso Scale Discovery (MSD) assay with a functional lower limit of quantification (LLOQ) of 144 pg/mL. NF-L was plotted against A) AAV dose, C) presence/absence of DRG toxicity (neuronal and/or nerve fiber degeneration), E) severity of DRG neuron degeneration/necrosis, and G) severity of DRG nerve fiber degeneration. Statistical significance ($p \leq 0.05$) was evaluated through one-way ANOVA analysis followed by Dunnett's multiple comparisons test (A, E, G) or t-test (C) performed in GraphPad Prism 9.0.0. NF-L for each animal has also been plotted by B) AAV dose, D) presence/absence of DRG toxicity (neuronal and/or nerve fiber degeneration), F) severity of DRG neuron degeneration/necrosis, and H) severity of DRG nerve fiber degeneration. Any values which fell below the limit of detection (LOD) were plotted as the functional LOD for that plate.

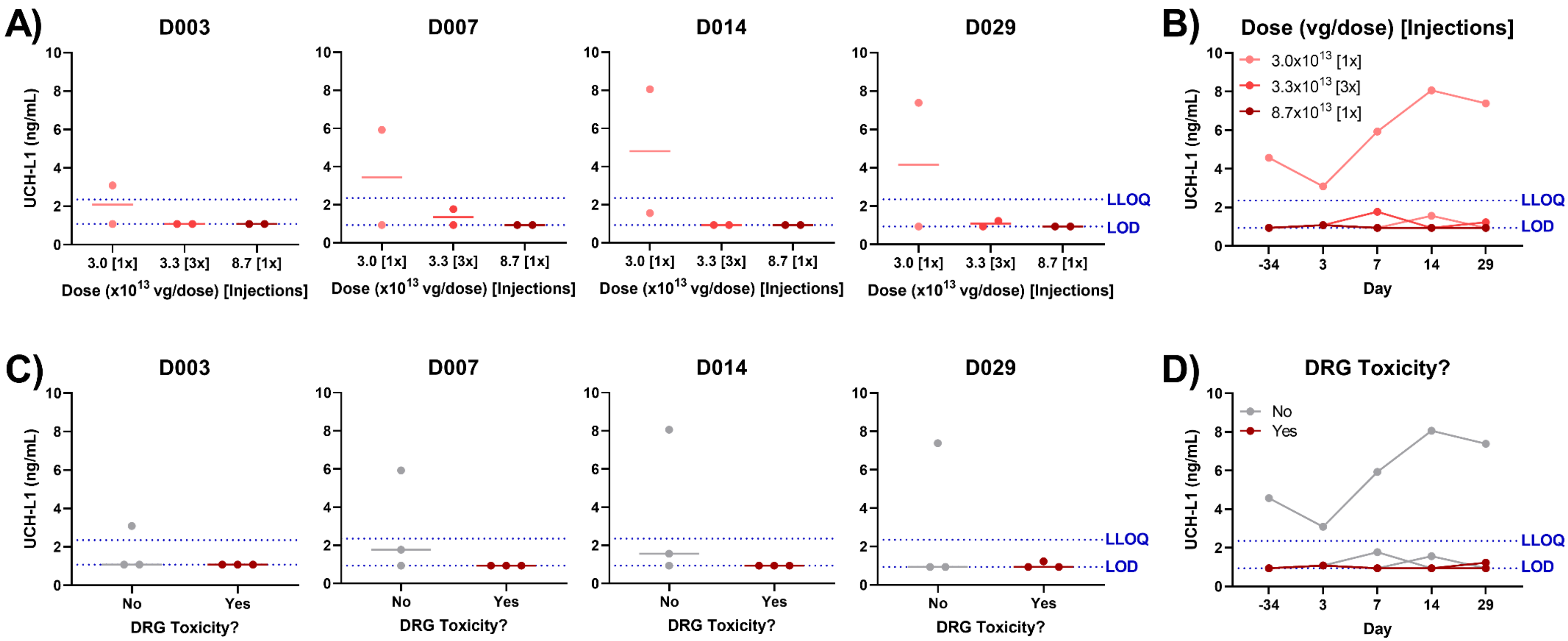


**Supplementary Figure S3: UCH-L1 in serum from cynomolgus monkeys exhibiting AAV-induced dorsal root ganglia (DRG) toxicity.** Male cynomolgus macaques (n=2 per group) were injected with $3.0x10^{13}$ [1x], $3.3x10^{13}$ [3x], or $8.7x10^{13}$ [1x] vg/dose AAV vector via intrathecal catheter. Serum was collected 3, 7, 14, and 29 days after injection, while histological DRG lesions were assessed on Day 29. UCH-L1 in serum was quantified using an in-house duplex Meso Scale Discovery (MSD) assay. The blue dashed lines indicate the limit of detect (LOD) and lower limit of quantification (LLOQ) for the assay. UCH-L1 was plotted against A) AAV dose and C) presence/absence of DRG toxicity (neuronal and/or nerve fiber degeneration). Statistical significance ($p \leq 0.05$) was evaluated through one-way ANOVA analysis followed by Dunnett's multiple comparisons test (A) or t-test (C) performed in GraphPad Prism 9.0.0. The UCH-L1 profile for each animal was also plotted by B) AAV dose and D) presence/absence of DRG toxicity (neuronal and/or nerve fiber degeneration). Any values which fell below the limit of detection (LOD) were plotted as the functional LOD for that plate.

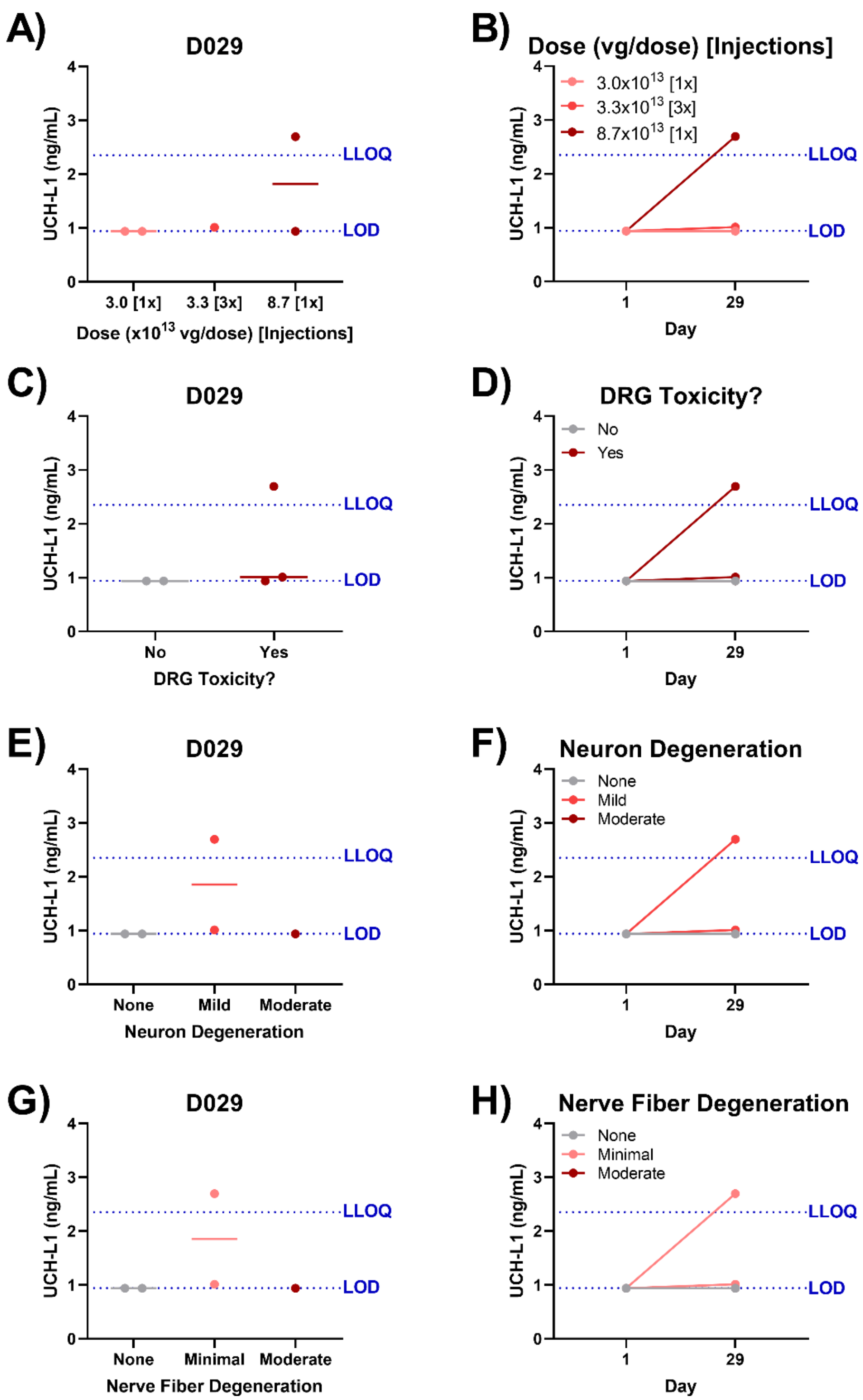


**Supplementary Figure S4: UCH-L1 in cerebrospinal fluid (CSF) from cynomolgus monkeys exhibiting AAV-induced dorsal root ganglia (DRG) toxicity.** Male cynomolgus macaques (n=2 per group) were injected with 3.0x$10^{13}$ [1x], 3.3x$10^{13}$ [3x], or 8.7x$10^{13}$ [1x] vg/dose AAV vector via intrathecal catheter. CSF was collected on Day 1 (pre-dose) and 29, while histological DRG lesions were assessed on Day 29. UCH-L1 in CSF was quantified using an in-house duplex Meso Scale Discovery (MSD) assay. The blue dashed lines indicate the limit of detect (LOD) and lower limit of quantification (LLOQ) for the assay. UCH-L1 was plotted against A) AAV dose, C) presence/absence of DRG toxicity (neuronal and/or nerve fiber degeneration), E) severity of DRG neuron degeneration/necrosis, and G) severity of DRG nerve fiber degeneration. Statistical significance ($p \leq 0.05$) was evaluated through one-way ANOVA analysis followed by Dunnett's multiple comparisons test (A, E, G) or t-test (C) performed in GraphPad Prism 9.0.0. The UCH-L1 profile for each animal was also plotted by B) AAV dose, D) presence/absence of DRG toxicity (neuronal and/or nerve fiber degeneration), F) severity of DRG neuron degeneration/necrosis, and H) severity of DRG nerve fiber degeneration. Any values which fell below the limit of detection (LOD) were plotted as the functional LOD for that plate.

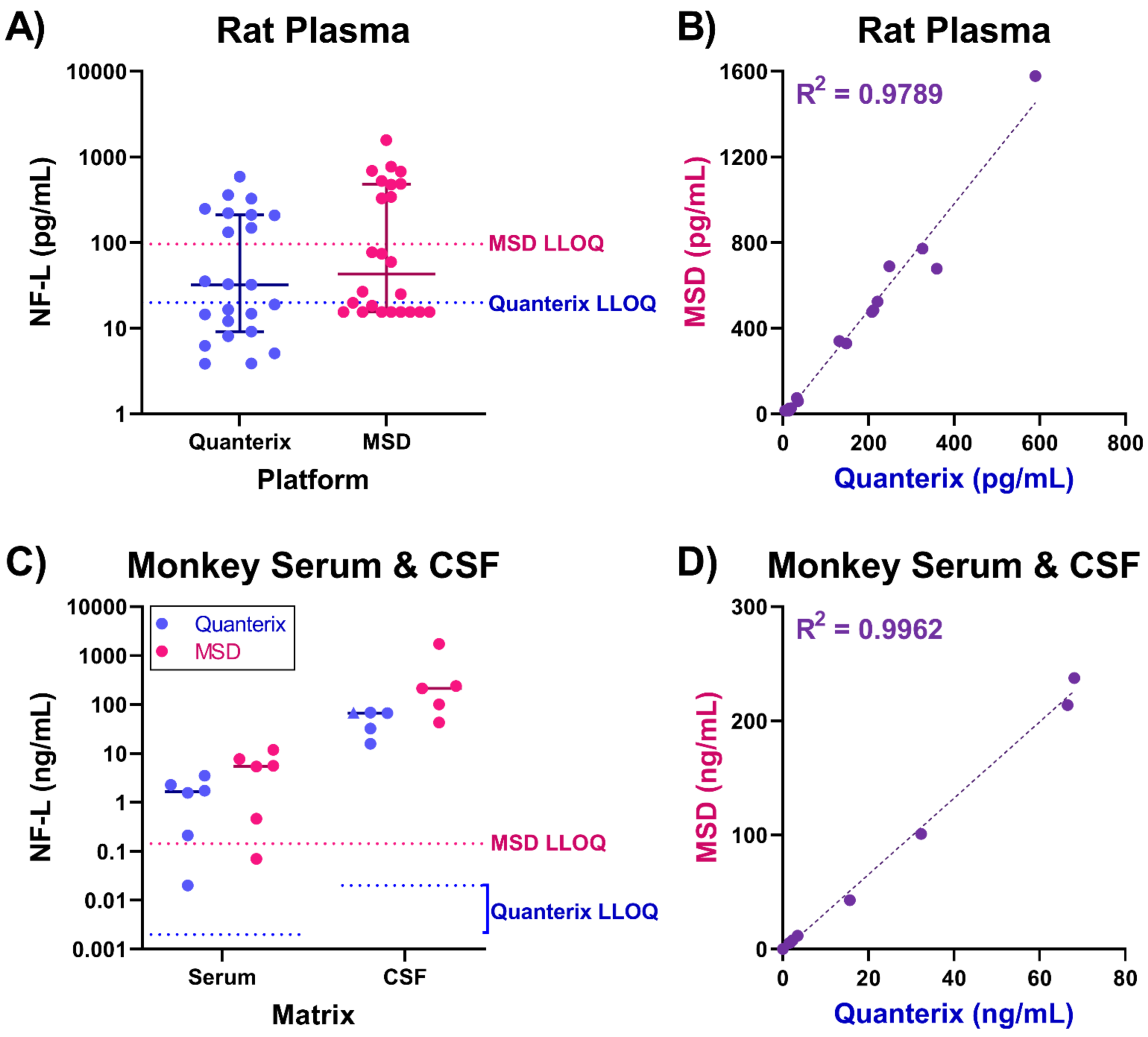


**Supplementary Figure S5: NF-L values detected with an in-house Meso Scale Discovery (MSD) assay are highly correlated with Quanterix Simoa HD-X values.** NF-L in A) rat plasma and C) cynomolgus monkey serum and cerebrospinal fluid (CSF) was evaluated using both the Quanterix Simoa Neurology 4-Plex Panel (blue) and an in-house duplex MSD assay (pink). The blue and pink dashed lines indicate the lower limit of quantification (LLOQ) for the Quanterix and MSD assays, respectively. The correlation between Quanterix and MSD values is shown for B) rats and D) monkeys, where the coefficient of determination ($R^2$) was determined through linear regression analysis. Any values which fell below the limit of detection (LOD) were plotted as the functional LOD for that assay. Note: The triangular data point (plot C) represents a sample which saturated the assay signal and thus has been plotted as the highest value obtained on the plate which did not saturate the signal. This sample was excluded from the correlation analysis (D).

### SUPPLEMENTARY TABLES

**Supplementary Table S1:** ‘Fit-for-purpose’ validation criteria

| Test | Replicates | Samples | Criteria |
|---|---|---|---|
| **INTRA-assay Precision & Accuracy** | 1 plate<br>≥ 3 replicates | **QC samples in matrix**<br>• High [ ]<br>• Mid [ ]<br>• Low [ ]<br><br>**Detection limits in diluent**<br>• ULOQ<br>• LLOQ | **For QCs:**<br>CV ≤ 25% between replicates<br>Bias = ± 25%<br><br>**For ULOQ & LLOQ:**<br>CV ≤ 25% between replicates<br>Bias = ± 30% |
| **INTER-assay Precision & Accuracy** | 3 plates<br>≥ 3 replicates | **QC samples in matrix**<br>• High [ ]<br>• Mid [ ]<br>• Low [ ]<br><br>**Detection limits in diluent**<br>• ULOQ<br>• LLOQ | **For QCs:**<br>CV ≤ 25% between plates<br>Bias = ± 25%<br><br>**For ULOQ & LLOQ:**<br>CV ≤ 25% between plates<br>Bias = ± 30% |
| **Freeze/Thaw Stability** | 1 plate<br>3 replicates | 1-4 freeze/thaw cycles of High, Mid, and Low QC samples | Recovery vs. Initial Thaw = 75-125% |
| **Dilution Linearity** | 1 plate<br>3 replicates | Serial dilution (2- to 64-fold) in matrix | $R^2$ ≥ 0.9 |

**Supplementary Table S2:** Sensitivity of Meso Scale Discovery (MSD) duplex assay

| Parameter | NF-L (pg/mL) | UCH-L1 (pg/mL) |
|---|---|---|
| Limit of Detection (LOD) | | |
| Analytical (average) | 7.37 | 213 |
| Functional (DF = 4) | 29.50 | 851 |
| Functional (DF=6) | 44.24 | 1,277 |
| Lower Limit of Quantification (LLOQ) | | |
| Analytical | 24 | 391 |
| Functional (DF = 4) | 96 | 1,564 |
| Functional (DF=6) | 144 | 2,346 |
| Upper Limit of Quantification (ULOQ) | | |
| Analytical | 25,000 | 400,000 |
| Functional (DF = 4) | 100,000 | 1,600,000 |
| Functional (DF=6) | 150,000 | 2,400,000 |

**Supplementary Table S3:** Assay validation results for rat and cynomolgus monkey biological matrices based on criteria listed in Supplementary Table S1

| | Rat | | | Cynomolgus Monkey | | |
|---|---|---|---|---|---|---|
| | **Plasma** | **Serum** | **CSF** | **Plasma** | **Serum** | **CSF** |
| **Validation Type** | Full (‘fit-for-purpose’) | Bridging | Bridging | Bridging | Bridging | Bridging |
| **Dilution Factor** | 4 | 4 | 6 | 6 | 6 | 6 |
| **INTRA-assay Precision & Accuracy** | Passed | NF-L Passed UCH-L1 Failed | Passed | NF-L Passed UCH-L1 Failed | NF-L Passed UCH-L1 Failed | Passed |
| **INTER-assay Precision & Accuracy** | Passed | Not evaluated | Not evaluated | Not evaluated | Not evaluated | Not evaluated |
| **Freeze/Thaw Stability** | Passed 4 cycles at -80ºC | Not evaluated | Not evaluated | Not evaluated | Not evaluated | Not evaluated |
| **Dilution Linearity** | Passed | Passed | Passed | Passed | Passed | Passed |
